# Discovery of the closest free-living relative of the domesticated “magic mushroom” *Psilocybe cubensis* in Africa

**DOI:** 10.1101/2024.12.03.626483

**Authors:** Alexander J Bradshaw, Cathy Sharp, Breyten Van Der Merwe, Keaton Tremble, Bryn T.M. Dentinger

## Abstract

The “magic mushroom” *Psilocybe cubensis* is cultivated worldwide for recreational and medicinal uses. Described initially from Cuba in 1904, there has been substantial debate about its origin and diversification. The prevailing view, first proposed by the *Psilocybe* expert Gastón Guzmán in 1983, is that *P. cubensis* was inadvertently introduced to the Americas when cattle were introduced to the continents from Africa and Europe (∼1500 CE), but that its progenitor was endemic to Africa. This hypothesis has never been tested. Here, we report the discovery of the closest wild relative of *P. cubensis* from sub-Saharan Africa, *P. ochraceocentrata* nom. prov. Using DNA sequences from type specimens of all known and accessable African species of *Psilocybe*, multi-locus phylogenetic and molecular clock analysis strongly support recognizing the African samples as a new species that last shared a common ancestor with *P. cubensis* ∼1.5 million years ago (∼710k - 2.55M years ago 95% HPD). Even at the latest estimated time of divergence, this long predates cattle domestication and the origin of modern humans. Both species are associated with herbivore dung, suggesting this habit likely predisposed *P. cubensis* to its present specialization on domesticated cattle dung. Ecological niche modeling using bioclimatic variables for global records of these species indicates historical presence across Africa, Asia, and the Americas over the last 3 million years. This discovery sheds light on the wild origins of domesticated *P. cubensis* and provides new genetic resources for research on psychedelic mushrooms.

## INTRODUCTION

*Psilocybe cubensis* (Earle) Singer is the most widely known, collected, and cultivated "magic mushroom" in the world (A. J. Bradshaw et al., 2022). *P. cubensis* was first described as *Stropharia cubensis* Earle from a cattle-grazed field in Cuba in 1904 and today is globally distributed where it is common in association with domesticated cattle across subtropical and tropical regions of America, Asia, and Australia (T. Froese et al., 2016; Guzman, 2005; McTaggart et al., 2023; Thomas et al., 2002). *P. cubensis* is one of the core species of psychoactive mushrooms used traditionally and contemporaneously for cultural and spiritual ceremonies across Mexico, and has been domesticated with many strains developed by an active social subculture due to its ease of cultivation (Castro Jauregui et al., 2022; Guzmán, 2008; Van Court et al., 2022). *P. cubensis* is also the target of ongoing biochemical and biomedical studies for drug discovery and whole organism therapies for a wide range of psychiatric illnesses (Blei et al., 2020; Brownstien et al., 2024; Fricke et al., 2017; Lerer et al., 2024; Matsushima et al., 2009; Shahar et al., 2024; Zhuk et al., 2015). Yet, despite the cultural, scientific and medical importance of *P. cubensis*, we know little of its specific ecology or evolutionary origins.

The prevailing hypothesis, first proposed by Guzmán (1983), is that *P. cubensis* originated in Africa and was transported to the Americas by Spanish colonizers during the 15th and 16th centuries. This hypothesis was largely predicated on the speculation that other species closely related to it remain to be discovered in Africa, a prediction rooted in the observation that the African continent is historically undersampled for *Psilocybe* (Piepenbring et al., 2020; Tsakem et al., 2024). The closest relatives of *P. cubensis* were recently shown to be from both Asia and Africa, but sampling from these and other regions remains vastly incomplete and definitive statements about origins and biogeography have remained untenable (A. J. Bradshaw et al., 2024). Interestingly, despite substantial popular knowledge and conspicuousness of *P. cubensis* in cattle-grazed pastures, it has not been officially confirmed from subtropical or tropical Africa, although casual reports of a *P. cubensis* lookalikes have appeared in peer-reviewed literature (Froese et al. 2016) and online databases such as iNaturalist and MycoPortal. *P. cubensis* does occur in Asia, recorded for Thailand (Ma et al. 2014) and India based on an ITS sequence matching *P. cubensis* in GenBank (accession OK165610.1), and multiple records of it occur across South and Southeast Asia in iNaturalist and MycoPortal.

One obstacle to determining the existence of named species in a given place at a given time is the availability of type specimens as references. Molecular identification is particularly sensitive to the lack of type reference sequences because sequences in public databases can carry names that have not been validated through comparison with them. In the absence of type reference sequences, this can result in a broad misapplication of names resulting in a collective misunderstanding of species’ identities, ecologies, and distributions. The recent study by Bradshaw et al. (2024) explicitly targeted type specimens for whole genome sequencing in an effort to remedy this long-standing problem in Psilocybe. While many type specimens were included in the study, it was not exhaustive, and critical types such as S. cubensis and species known only from Africa were not included.

Recent fieldwork across sub-Saharan Africa between 2013 and 2022 resulted in multiple specimens of an unknown Psilocybe sp. that is superficially similar to P. cubensis in habit, habitat, and general appearance. Upon further comparison of microscopic and molecular characters with the type of S. cubensis and other P. cubensis specimens from around the world, the recognition of the African specimens as a distinct species is warranted, provisionally named Psilocybe ochraceocentrata. A multi- locus molecular phylogenetic analysis was used to confirm its close relationship to P. cubensis, and divergence dating, ecological niche modeling, and species distribution modeling were used to predict when, in time, these two lineages diverged and where they were likely to be found over the last 3 million years.

## MATERIALS AND METHODS

### Collections and Sampling

Six collections of mushrooms resembling P. cubensis were collected across Zimbabwe and South Africa, occurring on or near decomposing herbivore dung. All collections were made on public or private land with the owner’s permission. Sporcarp collections were air-dried over low heat and preserved from insect damage using naphthalene. All other voucher details and deposition are reported in Table 1.

**Table 1:**
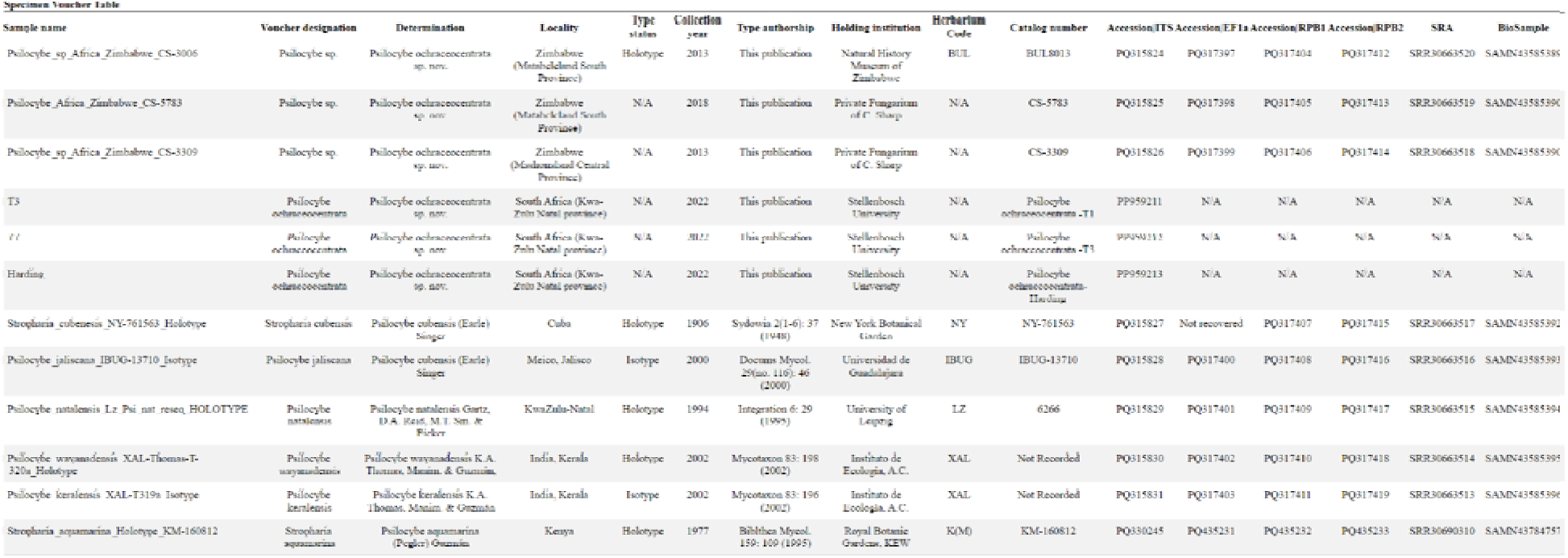
Sample and voucher information.

### Genomic sequencing, assembly, and barcode extraction from Type specimens

Type specimens of the Cubensae complex (Table 1) were extracted and sequenced in the same manner as (A. J. Bradshaw et al., 2024). Hymenophore fragments (5 to 15 mg) from dried fungarium samples were homogenized by placing them in 2.0 mL screw-cap tubes containing a single 3.0-mm and 8 × 1.5- mm stainless steel beads and shaking them in a BeadBug™ microtube homogenizer (Sigma-Aldrich, #Z763713) for 120 s at a speed setting of 3,500 rpm. DNA extraction of mechanically homogenized samples was performed using a Phenol-chloroform DNA extraction protocol. Lysis was performed with Monarch® Genomic DNA Purification Kit (NEB, #T3010S) following the manufacturer’s protocol for Tissue Lysis with an overnight incubation at 56 °C , using a volume of 500 ul lysis buffer, 10ul of Proteinase K and increasing the amount of wash buffer to 550 μL during each of the wash steps. Overnight incubation was then followed by a 2-hour incubation with 4ul of RNase A, after which total lysate was placed in homemade Phase Lock gel tubes made using Dow Corning™ High Vacuum Grease (ASIN:B001UHMNW0) along with an equal volume of OmniPur® Phenol:Chloroform:Isoamyl Alcohol (25:24:1, TE-saturated, pH 8.0) solution (MilliporeSigma, Calibiochem #D05686) and then mixed by gentle inversion for 15 min using a fixed speed tube rotator. After mixing, tubes were centrifuged at maximum speed (14,000×g) for 10 min; then, the aqueous (top) layer was transferred to a new phase-lock gel tube and the process repeated. DNA precipitation of the aqueous phase was performed by adding 5 M NaCl to a final concentration of 0.3 M and two volumes of room temperature absolute ethanol, inverting the tubes 20× for thorough mixing followed by an overnight incubation at −20 °C. The next day, DNA was pelleted by centrifugation at 14,000×g for 5 min. The DNA pellet was washed twice with freshly prepared, ice-cold 70% ethanol, air-dried for 15 min at room temperature, and then resuspended in 150 μL of Elution Buffer from the Monarch® Genomic DNA kit.

DNAs were then cleaned using the Zymo Research™ DNA Clean & Concentrator-5 (#D4003) kit, with the 5:1 binding buffer protocol to account for the short fragments common of older herbarium samples. Cleaned gDNAs were submitted to the High Throughput Genomics Core at the University of Utah, where sequencing libraries were prepared using the Nextera™ DNA Flex Library Prep (Illumina®, #20018704) and sequenced on a full lane of Illumina® NovaSeq 6000 PE 2 × 150 bp using an S4 flow cell. SRA accession and biosample numbers are provided in Table 1. Raw reads were trimmed using fastP v0.23.4 (Chen et al., 2018), assembled using MetaSPades v3.15.5 (Nurk et al., 2017; Prjibelski et al., 2020), and barcodes were extracted using Pathracer v3.16.0.dev (Shlemov & Korobeynikov, 2019) with custom hidden Markov models (A. Bradshaw et al., 2023). Genome assembly stats and resulting BUSCO (Simão et al., 2015) scores are reported in Supplementary Table 2.

### DNA barcoding and sequencing of additional Specimens

The Internal transcribed spacer (ITS) DNA barcodes, commonly used for species delineation (Schoch et al., 2012), of specimens “T1,” “T2,” and “Harding” were amplified using the primers set ITS1 and ITS4 (White et al., 1990). All ITS sequences were sequenced using Sanger sequencing and processed in the same manner as Van Der Merwe et al. (2024). gDNA from dried basidiomata was extracted using the Zymo Research™ Quick-DNA Fungal/BacterialMiniprep kits ( #D6005). Approximately 10 mg of fungal tissue was used for extraction, which was carried out according to the manufacturer’s instructions. Successful DNA extraction was visualized on a 1% agarose gel with ethidium bromide. Amplified barcodes were sequenced at the Central Analytical Facility, Stellenbosch University, using an ABI 3730xl DNA analyzer (Thermo Fisher Scientific™, Waltham, Massachusetts) with a 50 cm capillary array and POP-7. All ITS sequences were deposited into NCBI GenBank (Sayers et al., 2019) with accession numbers reported in Table 1.

### Phylogenetic Inference

Four independent loci were selected for phylogenetic analysis: the nuclear ribosomal internal transcribed spacers (ITS), translation elongation factor 1-alpha (Ef1a), the largest subunit of RNA polymerase II (RPB1), and the second largest subunit of RNA polymerase II (RPB2). In addition to newly generated sequences, selected sequences from NCBI GenBank (Sayers et al., 2019) and the UNITE database sh_general_release_dynamic_04.04.2024 (Abarenkov et al., 2010), including sequences from recently named species from Africa and elsewhere (Canan et al., 2024; Ostunii et al., 2024; Van Der Merwe et al., 2024) and sequences from type specimens (A. Bradshaw et al., 2023; A. J. Bradshaw et al., 2024); Supplementary data, Table 1), were used for phylogenetic analysis. ITS sequences were trimmed at the 5’ and 3’ end conserved motifs 5’- CATTA- and -GACCT-3’ for downstream analysis (Dentinger et al., 2010). Only the coding sequence of the protein-coding genes were used. ITS barcodes were partitioned into ITS1, 5.8S, and ITS2, using ITSx v1.1.3 (Bengtsson-Palme et al., 2013), aligned separately using MAFFT v7.490 with the flags --maxiterate 1000 --localpair for slow but accurate analysis (Katoh, 2002). Concatenation of multiple sequence alignments of the three ITS partitions was achieved using SEGUL 0.22.1 (Handika & Esselstyn, 2024). Phylogenetic analyses were performed using IQTREE2 2.3.6 with the flags -m MFP+MERGE -bnni -bb 1000, to enable automatic model finder (Kalyaanamoorthy et al., 2017) and 1000 ultrafast bootstrap replicates (Minh et al., 2013). Evolutionary models were automatically determined for ITS1, 5.8S, and ITS2 partitions separately, and for each codon position for EF1a, RPB1, and RPB2. Edges were linked across all partitions. Phylogenetic trees were rendered in FigTree (http://tree.bio.ed.ac.uk/software/figtree/).

### Molecular dating

A reduced dataset was constructed using sequences of the three protein-coding loci for all specimens represented by two or more loci. A concatenated matrix was constructed using Segul (Handika & Esselstyn, 2022), and each codon position was given its own partition. A root calibration normally distributed around a mean of 67.6 Ma (SD +- 6) following Bradshaw et al. (2024), and a calibrated Yule speciation model were used to estimate divergence times. Bayesian inference was conducted using BEAST2 v 2.7.7 (Bouckaert et al., 2019) with Markov Chain Monte Carlo (MCMC) sampling to estimate posterior distributions of phylogenetic parameters. The best fitting models of evolution determined with automatic model finder in IQTREE (Nguyen et al., 2015) were used for each partition. A chain length of 1 x 10^8^ steps was used, sampling every 5000 steps to ensure a thorough parameter space exploration. Convergence and effective sample sizes (ESS) of parameter estimates were assessed using Tracer v1.7.2 (Rambaut et al., 2018), with ESS values above 200 considered indicative of sufficient mixing and convergence. The resulting posterior tree distribution was downsampled to every 2000 trees using LogCombiner due to computational limitations and then summarized using TreeAnnotator v1.10 (Drummond & Rambaut, 2007) to produce a maximum clade credibility tree, with node heights representing median posterior estimates. The annotated tree was visualized using FigTree as above, with posterior probabilities indicated on nodes to reflect the robustness of inferred relationships and 95% highest posterior density (HPD) of estimated divergence times indicated with bars.

### Species Distribution Modeling

To predict the ranges of *P. cubensis* and *P. ochraceocentrata*, we conducted species distribution modeling (SDM) using GPS coordinates of known collections and the contemporary 19 bioclimatic variables at 2.5M resolution (Fick & Hijmans, 2017) as environmental predictors. In addition, we sought to predict their historical ranges by conducting SDM of *P. cubensis* using four 19 bioclimatic datasets modeled to have occurred during the Current age, Anthropocene (1979 – 2013), last interglacial (LIG) ∼130KYA , Pleistocene, MIS19 (∼787 KYA) and the Pliocene ∼3.3Mya (Brown et al., 2018; Dolan et al., 2015; Hill, 2015; Karger et al., 2017) at 2.5M resolution accessed from paleoclim.com (Brown et al., 2018). All collection GPS coordinates used for SDM were pulled from MycoPortal (Miller & Bates, 2017) entries, including data from Mushroom Observer (https://mushroomobserver.org/) and iNaturalist (https://www.inaturalist.org). All data points were filtered to remove non-wild collections, including the removal of all entries with specific mentions of samples being cultivated, confiscated by police, or those labeled as known cultivated strains of *P. cubensis*. The locations of all collections and observations were plotted using GPS coordinates over a global map with R packages ggmap v. 4.0.0 (Kahle & Wickham, 2013) and sf v1.0-16 (Pebesma, 2018), and then visualized using ggplot2 v3.5.1 (Wickham, 2016).

To account for lumping of *P. ochraceocentrata* in prior records of *P. cubensis*, SDM for all environmental datasets was conducted both with and without African collections of *P .cubensis*, using the SDM R package (v1.2-46). We tested six of the most common SDM models ("bioclim", "domain.dismo", "glm", "gam", "rf", and "svm"), using 1000 random points as “absence” points to validate the model (Supplement Data). The best-performing model, according to AUC, COR Deviance, TSS, MCC, and F1 score, was chosen for each environmental dataset.

### Microscopy and Morphological analysis

Microscopic analysis on specimens CS-3006, CS-5783, and CS-3309 was performed using a Leitz Wetzlar Orthoplan microscope with a Leitz Wetzlar drawing tube. All spore measurements were taken from thirty spores whose side-view was clearly visible. Spore prints were studied using Melzer’s Reagent, and sections of dried lamellae were studied in Congo Red. Color descriptions were derived from (Rayner, 1970), and codes are recorded parenthetically for each specimen. Microscopic analysis of additional specimens T1, T3, and Harding was performed using Methods outlined in Van Der Merwe et al. (2024). The length and width were measured, and the length/width quotient (Q) was calculated and reported in Supplementary Table 1 for all samples.

## RESULTS

### Molecular systematics

Independent phylogenetic analysis of ITS, EF1a, and RPB2 all recovered African specimens originally identified as *P. cubensis*, *P.* cf. *cubensis*, and *P.* cf. *natalensis* as a monophyletic group sister to a clade containing sequences from the holotype of *Stropharia cubensis* Earle (=*Psilocybe cubensis*) (Fig. 1) with strong bootstrap support (ITS=97%, EF1a=99%, RPB2=100%). These same African sequences were not reciprocally monophyletic with respect to sequences of *P. cubensis* using RPB1. Sequences of the types of the other known species of *Psilocybe* endemic to Africa were consistently recovered in other clades. Sequences from the holotype of *Stropharia aquamarina* Pegler were placed sister to *Psilocybe wayanadensis* K.A. Thomas, Manim. & Guzmán and other species from Asia and Australia (BS; ITS=100, EF1a=92, RPB1=88, RPB2=97, Concat=100). Sequences from the holotype of *Psilocybe natalensis* Gartz, D.A. Reid, M.T. Sm. & Eicker did not match any publicly available sequences and was placed sister to *Psilocybe chuxiongensis* T. Ma & K.D. Hyde (typified from Yunnan, China) and *Psilocybe maluti* B. Van der Merwe, Rockefeller & K. Jacobs (typified from South Africa) ( BS; ITS=99, Concat=99). Sequences of the holotype of *Psilocybe jaliscana* Guzmán and the sequences of *Psilocybe subcubensis* Guzmán were nested within a clade of sequences from *P. cubensis* across all loci. Sequences from the holotype of *Psilocybe keralensis* K.A. Thomas, Manim. & Guzmán was recovered in Clade I, and its ITS sequence was identical to the ITS from the type specimen of *Psilocybe ingeli* B. Van der Merwe, Rockefeller & K. Jacobs.

**Figure 1.**
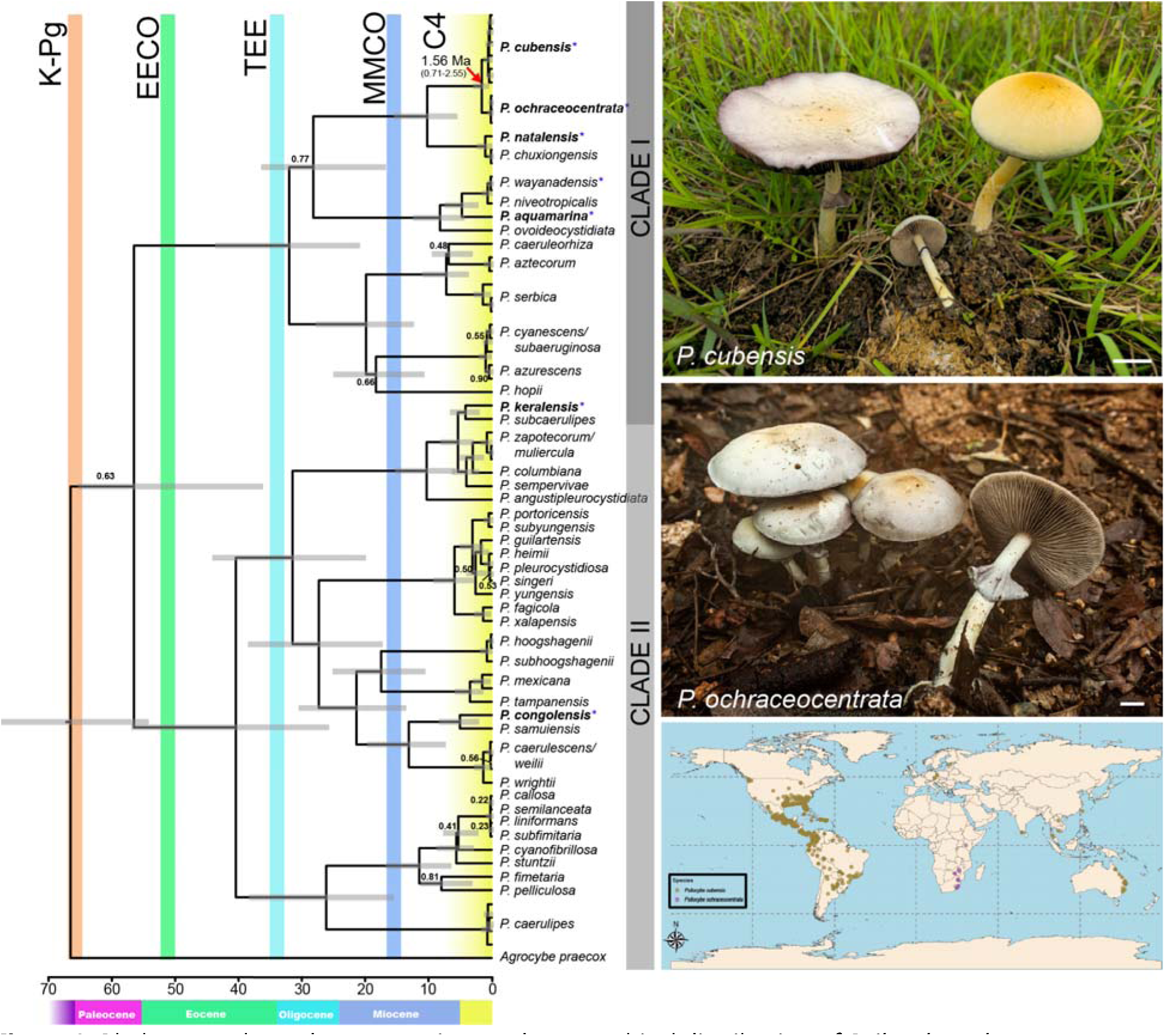
Phylogeny, photo documentation, and geographical distribution of Psilocybe ochraceocentrata. LEFT: Maximum clade credibility chronogram of Psilocybe spp. resulting from Bayesian divergence analysis of three loci (EF1a, RPB1, RPB2). All nodes received >95% posterior probability except for internal nodes with posterior probabilities indicated. Shaded bars at nodes represent 95%HPD. Shaded bars have been removed from nodes where descendants represent multiple individuals of a species for improved readability. The mean divergence time for the common ancestor of P. cubensis and P. ochraceocentrata is indicated by a red arrow. Scale bar at the bottom represents the estimated time using a root calibration normally distributed around a mean of 67.6 Ma (SD +- 6) following Bradshaw et al. (2024). Geologic epoch time scale is approximate. Colored vertical bars represent noteworthy global events that illustrate correlation of major divergences within Psilocybe: the K-Pg event ∼65 MYA, the Eocene Epoch Climate Optimum (EECO), the Terminal Eocene Event (TEE), the Mid-Miocene Climate Optimum (MMCO), and the emergence of C4 grass biomes (C4). Right: in situ photos of Psilocybe cubensis (BD1406, top), P. ochraceocentrata (CS3006, middle), and records of P. cubensis (gold squares) from Mycoportal, iNatrualist, and Mushroom Observer with African records reinterpreted as P. ochraceocentratta (Purple squares).

### Molecular dating and geological timeline

Due to the close relationship between *P. ochraceocentrata* and *P. cubensis*, we performed molecular dating using Bayesian inference with BEAST of a concatenated dataset of EF1a, RPB1, and RPB2 (Figure 1). Phylogenetic reconstruction generated a tree with highly supported nodes (posterior probability >95%). Only a minority of nodes received less than a 0.95 posterior probability and most of these were near the tips among closely related species and species complexes. Molecular dating places the MRCA of *P. ochraceocentrata* and *P. cubensis* at ∼1.56 million years ago (MYA)(0.71-2.55 95% HPD). This estimated divergence date corresponds to the Pleistocene epoch (2.5 MYA - 11.7 KYA) following the mass emergence of grass biomes in warm climates with the evolution of the C_4_ photosynthetic pathway (8 to 3 MYA) (Edwards et al., 2010).

### Ecological niche modeling and species distribution of P. cubensis

Due to the close relatedness of *P. ochraceocentrata* and *P. cubensis*, we chose to investigate the theoretical species distribution of *P. cubensis* through time using ecological niche modeling (ENM) and species distribution modeling (SDM). To do so, we used publicly available data from MycoPortal, which includes data from iNaturalist and Mushroom Observer. Occurrence data was filtered for only those with georeferences and removed any entries that suggested occurrences were acquired from non-natural sources, such as cultivation. After filtering, we found 1001 occurrences, 12 of which were from the African continent (Supplementary data). Due to the inability to authenticate specimens from Africa as *P. cubensis*, and sparse literature reports of its distribution across Africa and India (Guzmán, 2014; Thomas & Manimohan, 2003) , we performed ENM and SNM in two ways. The first analysis included all geopoints (minus confirmed *P. ochraceocentrata* specimens), assuming that *P. cubensis* specimens from Africa were correctly identified (Figure 2). The second analysis removed African specimens, assuming that these specimens are *P. ochraceocentrata* (Supplementary Figure 5). ENM and SDM were done using multiple geological datasets, ranging from the Pliocene (∼3MYA) to the Modern day (Figure 2, Supplementary data).

**Figure 2.**
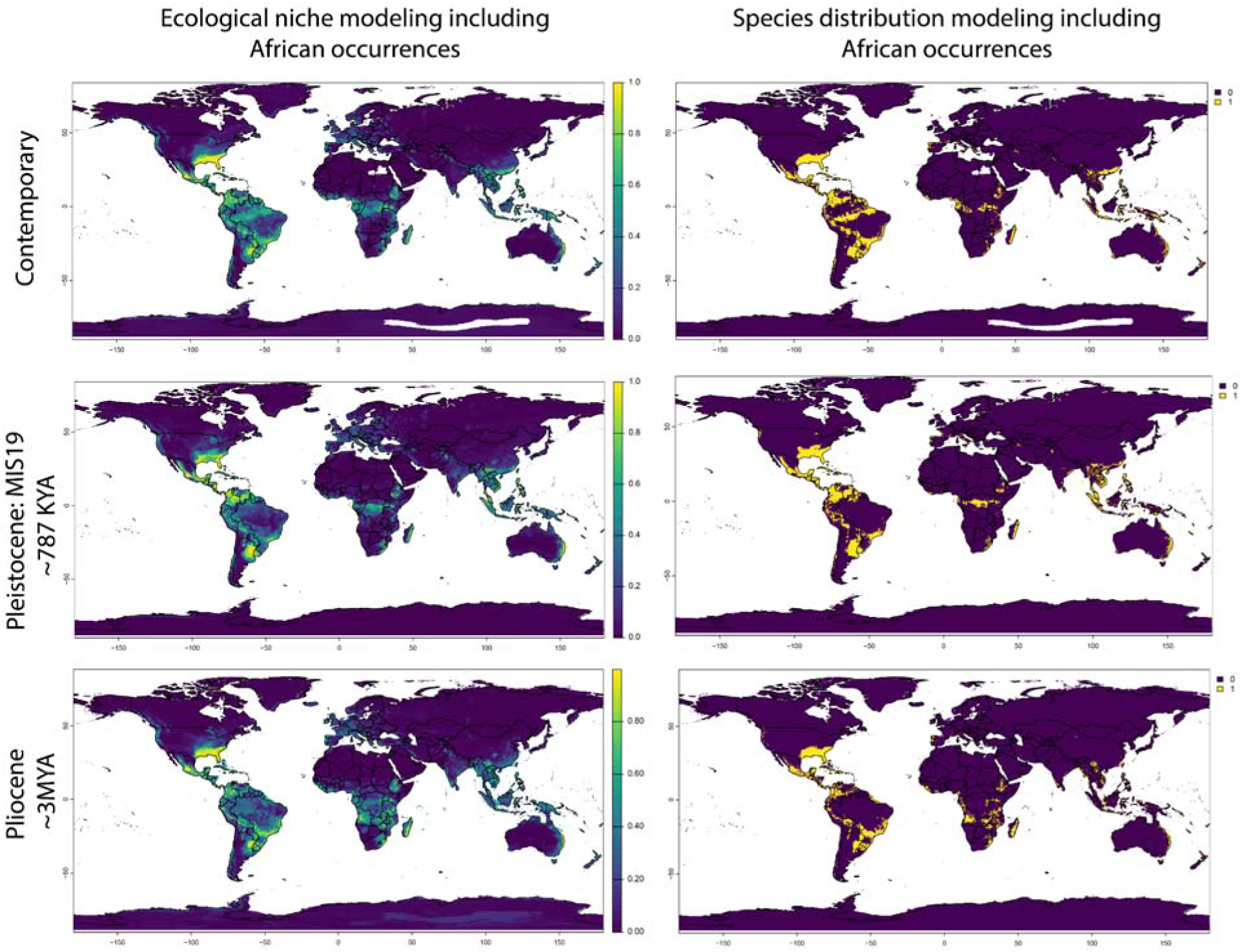
Ecological niche modeling (ENM) and Species distribution modeling (SDM) of Psilocybe cubensi and P. ochraceocentrata through time. All occurrence data for P. cubensis from public sources (likely including P. ochraceocentrata misidentified as P. cubensis in Africa) across three time scales since the mid-Pleistocene. **Top Row**: Contemporary (Modern day), **Middle Row**: Pleistocene, MIS19 (∼787 KYA) , **Bottom Row**: Pliocene (∼3MYA). **Left**: ENM with distribution likelihood indicated as a heat gradient from purple (0%) to yellow (100%). **Right**: SDM with predicted species presence as present (1, yellow) or absent (0, purple).

**Figure 3.**
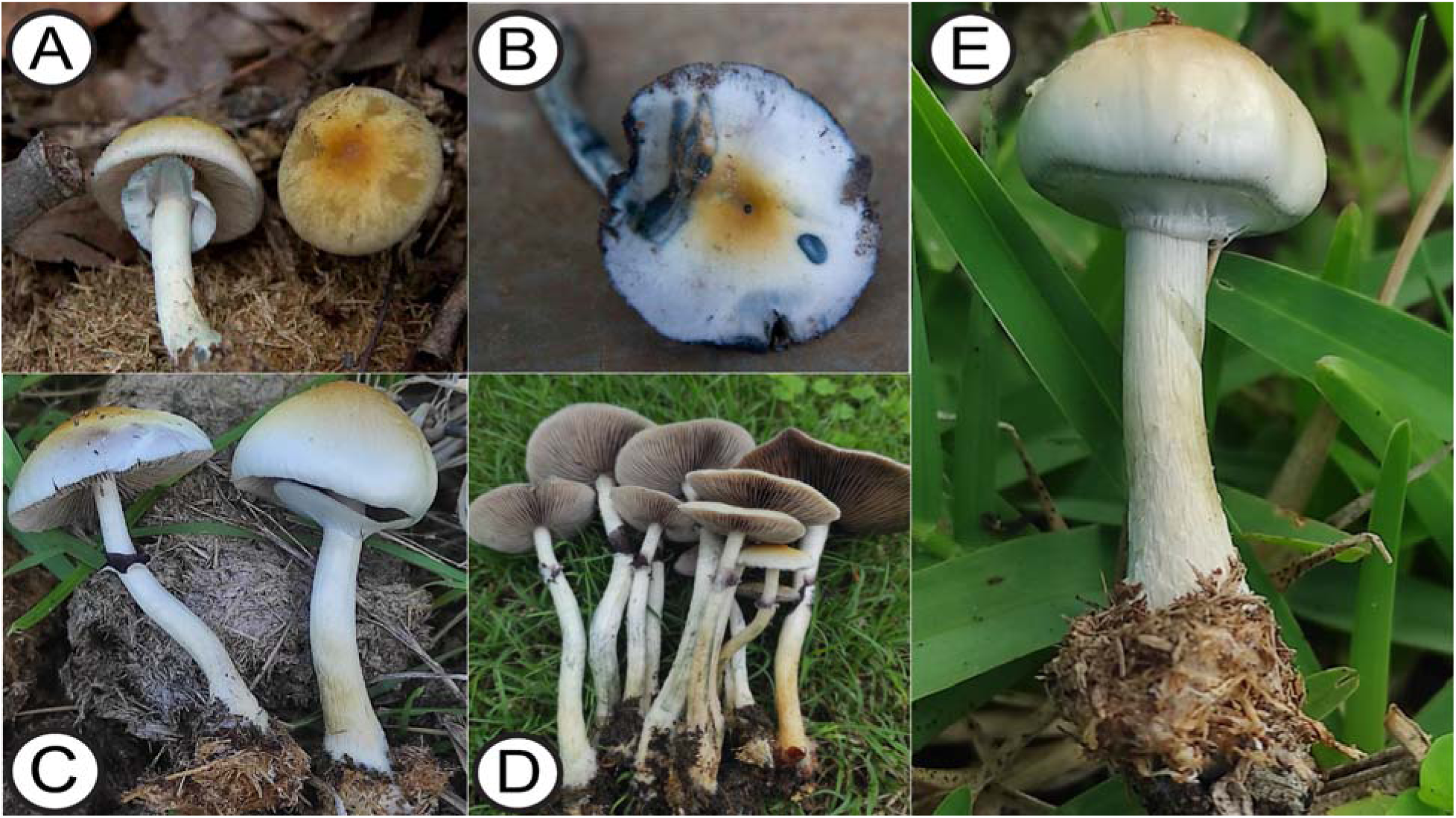
(A) *Psilocybe ochraceocentrata*. Holotype collection (CS-3006). Photo credit: C.Sharp,(B) Older fruiting body of *Psilocybe ochraceocentrata*; color change possibly due to water logging, (C)*Psilocybe ochraceocentrata* found on dung; (D) Cluster of *Psilocybe ochraceocentrata* (South Africa). Photo credit: Talan Moult, (E) Young sporocarp of *Psilocybe ochraceocentrata* (Harding). Photo credit: Talan Moult.

**Figure 4.**
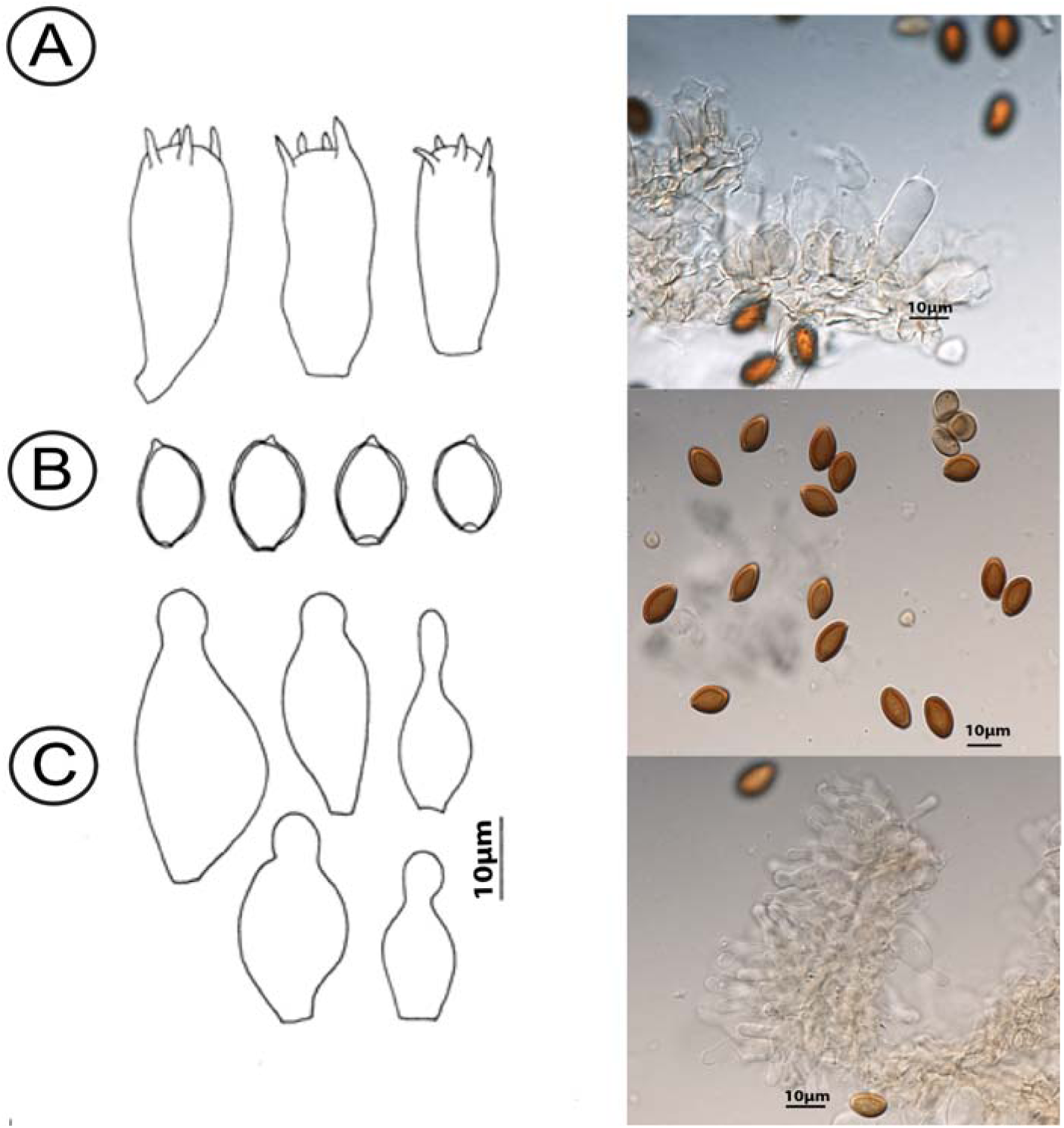
Microscopic illustrations and microscopic feature morphology of Psilocybe ochraceocentrata: (A)Basidia from Holotype specimen CS-3006; (B): Basidiospores from Holotype specimen CS-3006; (C) Cheilocystidia from specimen CS-5783. Microscopic images are derived from specimen P. ochraceocentrata "Harding.”

Both data sets exhibited high similarity with ENM and SDM across time, predicting similar species distributions (Figure 2, Supplementary Figure 5). The most extensive predicted ranges occurred across the Southern United States, Central America, and South America (ENM >0.8, SDM =1), with a presence across Southern Africa, Southeast Asia, and Australasia, albeit as a more restricted range (ENM<0.5, SNM=1) (Figure 2). Across time, ENM and SDM indicated reduced distribution across Africa, while the Americas, South East Asia, Australasia, and Europe exhibited an increased presence and distribution up to the Last Interglacial (LIG) ∼130 thousand years ago (KYA). Post LIG into contemporary times, ENM and SDM predict distributions more in line with the Pleistocene, MIS19 (∼787 KYA) and the Pliocene (∼3MYA), but with a slightly increased predicted presence in Africa and distribution across Southeast Asia and Australasia remaining similar to the LIG (Figure 2).

### Taxonomy

**Psilocybe ochraceocentrata** C. Sharp, A. Bradshaw, B. Dentinger & B. van der Merwe nov. sp. Index Fungorum Registration #: IF 902894

Holotype Deposition: Natural History Museum of Zimbabwe, BUL8013

GenBank Accession: PQ315824 (ITS), PQ317397( EF1a), PQ317404 (RPB1), PQ317412(RPB2)

SRA Accession:

### Etymology

Pileus with a yellow-ochre center.

### Diagnosis

Typus: Zimbabwe, Matabeleland South Province, The Farmhouse, Kezi Road, Matobo Hills. QDS 2028A4. In thick leaf litter in mixed deciduous woodland on granitic sand. 14 Jan 2013. Collectors C. Sharp & R. Aldridge. Holotype: CS3006 (Field ID, Private Fungarium of C. Sharp), Holotype split: Natural History Museum of Zimbabwe, BUL 8013.

### General field description

**Fruiting body** medium-sized, up to 105 mm tall and growing in tight clusters; a pale fungus that bruises blue-green. **Pileus** 65-75 mm diam.; first cream-colored, then vinaceous-grey (115) with center ochreous (44) to fulvous (43); pale yellow towards margin; convex then planate, often undulating; surface finely and radially silky with a dull sheen. **Flesh** bright cream-coloured, firm to pithy. **Margin** first down-curved; edge smooth often with radial cracking. **Lamellae** adnate; first pale then greyish-sepia (106) to brown- vinaceous (84) to very dark sepia and almost black near margin; face of gill speckled as spores ripen; to 12 mm deep, thin, papery and fragile; edge thin, smooth or finely scalloped, pale; sparsely to moderately crowded with lamellulae, 6-8/cm. **Stipe** central; up to 95 mm long x 9-10 mm; cream-coloured, bluing where handled; cylindrical, often twisted; apex minutely tufted (x10), streaked longitudinally, dull sheen, base often with white silky hairs. **Flesh** fibrous in walls and center hollow. **Ring** median to higher; white first then with dark spores; membranous, striate, very fragile and soon clinging to stipe and disintegrating. Mycelium is white to cream-coloured, forming a thick, compact mat amongst leaf litter. Bruising instantly to blue-green (94) where handled or damaged. **Odour** of typical mushroom. **Spore-print colour** dark vinaceous-grey (116) to purplish-grey (128).

### Habit and habitat

Miombo woodland, mixed deciduous woodland, all on granitic sand; CS3309 was found growing on old, decomposed herbivore dung of unknown origin.

### Macroscopic Description

**Fruiting body** in a tight cluster, to 105 mm tall. **Pileus** to 65 mm diameter; first cream-colored to pale vinaceous-grey (115) with ochreous (44) or fulvous (43) center; first convex then planate; surface finely streaked radially with dull sheen. **Flesh** cream-coloured, firm. **Margin** straw-yellow (40), down-curved, smooth; some radial cracking. **Lamellae** greyish-sepia (106) to brown vinaceous (84), face speckled with ripening spores; adnate; thin, papery, fragile; edge finely scalloped and pale; with lamellulae. **Stipe** to 95 mm long x 9 (apex) – 10 (mid) – 10 (base) mm; cylindrical, twisted or not; apex minutely tufted (x10 lens), longitudinally streaked to silky-hairy at the base. **Flesh** hollow with fibrous walls. **Ring** just above median, white, striate, membranous to clinging, and very fragile. **Bruising** immediately to glaucous sky-blue (93) to glaucous blue-green (94). **Odour** typical mushroom. **Spore-print** dark vinaceous-grey (116) to purplish-grey (128). **Chemical reactions:** not recorded

### Microscopic description

**Spores derived from Holotype CS-3006**,: Ranges are given parenthetically with mean values underlined; ellipsoid, a few lenticulars; thick-walled, smooth-walled with germ-pore; (11)<u>11.8</u>(12.5) x (6.5)<u>7.6</u>(8.0) µm; Q = (1.44)<u>1.55</u>(1.71). Basidia elongate or clavate and variable in size, 22-38 x 8-12.5 µm; four sterigmata, 4-5µm long with rounded or acute apex.

Notes from additionally examined samples:

1. [1] Spores can be highly variable in size; note CS3309 has much smaller spores with a few in the extreme range (Supplementary Table 1)
2. [2] **Spores** shape ellipsoid, some collections having more lens-shaped spores than others; thick-walled, smooth-walled with apical germ-pore. **Basidia** elongate or clavate; 22-30 x 8-11 µm; four sterigmata, 4- 5µm long with rounded or acute apex. **Cystidia** no pleurocystidia were observed; despite numerous sections, no cheilocystidia were observed in the Type collection but very clear in CS3309 and CS5783; capitate in shape, thin-walled, 21-26 x 9-13µm across the widest part. **Hyphae** clamp connections observed in the Type.
3. [3] CS-2680. Mashonaland Central Province, Mukuvisi Woodlands, Harare. QDS 1731C3. 18 Jan 2012. Collector Daniel Nyamajiwa. The spore range is a good match even if a bit smaller, e.g. (10)<u>10.8</u>(11.5)[13] x [6.5](7)<u>7.4</u>(8) µm but shape slightly different, Q = (1.33)<u>1.45</u>(1.57) (Supplementary Table 1).
4. [4] Even though the ochre-yellow center is diagnostic, one noticeable morphological difference is that the green-blue pigments turn black and remain dark while the specimen is fresh. This may be due to the older fruiting bodies or because they contain more water in a very wet season. All signs of green or black coloration are absent in the dried samples (Figure 2, B).

### Synonomizations

Psilocybe cubensis (Earle) Singer, Sydowia 2(1-6): 37 (1948)

Basionym:

*Stropharia cubensis* Earle, Inf. an. Estac. Cent. agr. Cuba 1: 240 (1906)

Synonyms:

=*Naematoloma caerulescens* Pat., Bull. Soc. mycol. Fr. 23(2): 78 (1907)

c*Hypholoma caerulescens* (Pat.) Sacc. & Trotter, Syll. fung. (Abellini) 21: 212 (1912) c*Psilocybe cubensis* var. *caerulescens* (Pat.) Singer & A.H. Sm., Mycologia 50(2): 269 (1958)

=*Stropharia cyanescens* Murrill, Mycologia 33(3): 279 (1941)

c *cubensis* var. *cyanescens* (Murrill) Singer & A.H. Sm., Mycologia 50(2): 269 (1958)

=*Psilocybe jaliscana* Guzmán, Docums Mycol. 29(no. 116): 46 (2000)

## DISCUSSION

*Psilocybe* has become a world-renowned genus of mushrooms, primarily due to their psychoactivity. Currently, *Psilocybe* mushrooms have been targeted as a source of natural products for use in a global mental health crisis (Carhart-Harris et al., 2017; Daniel & Haberman, 2017; Gandy et al., 2020; Johnson & Griffiths, 2017). However, due to the issue of legality in their collection, characterization of their diversity and studies of their biology have been severely stifled. Despite the ∼160 species of true *Psilocybe* described around the world, most were described in the Americas (Borovička et al., 2011; Guzmán, 2014; Guzman et al., 2013; Johnston & Buchanan, 1995; Ma et al., 2014; Picker & Rickards, 1970). In contrast, only seven species of *Psilocybe* (including *P. ochraceocentrata)* have been typified from Africa, ranging from the cedar forests of Northern Africa (*Psilocybe mairei)* to the grassland of South Africa (*Psilocybe maluti*) (Guzmán, 2014; Van Der Merwe et al., 2024); many more still undocumented species are predicted.

The subject of psychoactive mushrooms, in particular *Psilocybe*, and their proposed medical benefits is rapidly gaining interest globally, and no less so in Zimbabwe. Recreational use in Zimbabwe is generally isolated and often dependent on the availability of imported products, likely the frequently cultivated *P. cubensis* (Musshoff et al., 2000). However, historical indigenous knowledge of these fungi is lacking outside of Mesoamerica, making traditional-use claims challenging to validate. However, there have been suggestions that access to the information has been restricted due to a sense of protected information among traditional healers (T. Froese et al., 2016; Guzmán, 2014; Van Der Merwe et al., 2024). This issue is likely a consequence of the historical lack of mycological studies across Africa, which remains one of the most understudied geographic locals for fungal diversity (Antonelli et al., 2024; Crous et al., 2006). Consequently, the people of Zimbabwe’s use of *Psilocybe* for ceremonial or medicinal purposes is unknown.

*Psilocybe* has been previously grouped into four major sections, including the “Cordisporae”, “Mexicanae,” “Zapotecorum,” and “Cubensae” sections. Of these sections, the Cubensae has had little documentation of novel species. The Cubensae section includes species found across Central America, Southeast Asia, India, and Africa, including the most iconic species *P. cubensis* (A. J. Bradshaw et al., 2024; Guzmán, 1983, 1995; Ramírez-Cruz et al., 2013). However, the abundance of diversity within the section remains uncertain, primarily due to a limited representation of type specimens. Type specimens serve as the authoritative description of a species, and their representation is essential to characterize new species definitively. In particular, some specimens of the Cubensae complex, such as *Psilocybe jaliscana* (typified from Mexico) and *Psilocybe aquamarina* (typified from Kenya), were thought to be synonymous with *P. cubensis* and *P. subcubensis* , respectively (Guzman 2014, and personal Communication, Virginia Ramírez-Cruz and Alonso Cortés-Pérez). While the case for *Psilocybe jaliscana* being synonymous with *P. cubensis* is true, the same is not true for *P. aquamarina* , which is reported here as genuinely novel.

Further, a new issue arises when publicly deposited data with type specimens is validated. The commercially sold “Natal Super Strength (NSS)” (OK491080.1) strain of *P. natalensis* (typified from KwaZulu-Natal) does not match the type specimen of *P. natalensis* . Instead, four of the five publicly deposited sequences cluster with *P. ochraceocentrata*, indicating misidentification. This could lead to future regulatory issues and confusion over the species identity of commercially sold *Psilocybe* strains. Misidentifications at these levels illustrate the difficulty of identifying many species of *Psilocybe*, which often exhibit plastic morphology, necessitating type specimen validation for taxonomic accuracy and stability. This phenomenon is not unique to the Cubensae complex, as it has also been shown to occur across the genus multiple times in commonly collected species (Awan et al., 2018; A. J. Bradshaw et al., 2022, 2024). Without the ability to accurately describe what species are being consumed medically or being commercialized, we jeopardize future work in understanding the species-specific secondary metabolism of *Psilocybe.* With this work, we explicitly targeted type specimens, allowing for direct and accurate comparison of future specimens of *Psilocybe* from around the world. Notably, we provide the molecular data from the holotype of *Stropharia cubensis* Earle, confirming the identity of the majority of commercially available strains of “magic mushrooms” as *P. cubensis* for the first time.

*Psilocybe cubensis* is a globally distributed species whose origin is debated. Our ecological modeling suggests that *P. cubensis* could have been found historically but discontinuosly across the Americas, Southeast Asia, Southern and Central Africa, and Australasia between 0.71 and 2.55MYA (Figure 2, Supplementary data). While there is an extensive predicted range, it has been shown that *P. cubensis* from Australia has relatively low genetic diversity, suggesting recent arrival with domesticated cattle to the continent (McTaggart et al., 2023). There are also no authenticated specimens of *P. cubensis* from Africa and only two sequences of it from India (iNat144581118, OK165610.1). While it is possible that *P. cubensis* may occur naturally in Africa, *P. ochraceocentrata* has a strikingly morphological and genetic similarity and shares its almost exclusively ruminant coprophilic habit, which is likely to lead to misidentification. Our divergence analysis suggests that their MRCA likely originated alongside the large herbivores, possibly during the expansion of the C4 grasslands in East Africa 1.8-1.2 MYA (Cerling, 1992; Cerling et al., 1988). Coincidentally, this is also the period when *Homo erectus* became the dominant hominin in East Africa and the first to spread from Africa through Eurasia via the Levantine corridor alongside large herbivores, including bovids (Antón & Swisher, Iii, 2004; Belmaker, 2010; Dennell & Roebroeks, 2005; Zhu et al., 2018). These major migration events present a possible avenue for dispersal of the MRCA of *P. cubensis* and *P. ochraceocentrata* from Africa and their subsequent divergence in Asia and Africa after aridification lead to loss of habitat in the intervening region.

*P. cubensis* was typified from Cuba in 1904 and is regularly found across the Americas, where it is associated with herbivore dung. However, cattle did not reach the Americas until 1493 (Sluyter, 2023) during the European colonization of Mesoamerica. While the origins and modern domestication of cattle are still debated, there is no question that their evolutionary origins reside in the Old World as opposed to the New World. Domestication of large grazing cattle (*Bos taurus*) and Asian zebu (*Bos indicus*) occurred between ∼8-10KYA. The MRCA of both of these domesticated animals, aurochs (*Bos primigenius*), occupied a range from northern Africa to both coasts of Eurasia (Hanotte et al., 2002; Pitt et al., 2019; Zeuner, 1963), overlapping with estimated historic species distribution of *P. cubensis*.

Precolonial presence of *P. cubensis* in the Americas is not known, but its estimated divergence from *P. ochraceocentrata* and ecological niche modeling suggest it may have existed there before the arrival of Europeans. Bison (*Bison* spp.) are bovids that dominated grasslands in the Americas throughout the Pleistocene and may have facilitated the precolonial arrival and spread of *P. cubensis*. Bison are thought to have arrived from Asia to the Americas in multiple waves, with the first occurring ∼195–135KYA during the LIG and then the secondc45–21KYA (D. Froese et al., 2017). After their arrival, bison became one of the dominant herbivores in the Americas and were widely distributed across North America and Central America. Bison are known ecological engineers, expanding and maintaining grassland ecosystems wherever they roam (Gates et al., 2010; Ripple et al., 2015), making them important vectors for coprophilic fungi, potentially including *P. cubensis*. The predicted ranges that we see in the ENM and SDM results overlap with reported range of bison across the Americas, including their co-occurrence across the coastlines of Alaska and the Pacific Northwest, and the peninsula of Florida during the LIG (Supplementary data). The occurrence of *P. cubensis* in the Antilles archipelago is most likely a recent development since no large herbivore existed that could have supported it except for giant ground sloths (*Megalocnus* spp.), although this seems unlikely given these were probably browsers with grasses making up a limited portion of their diets (Dantas et al., 2023). While we cannot confirm the Americas are an origin of *P. cubensis*, we also cannot rule it out with the presently available data. However, the inferred divergence of *P. cubensis* and *P. ochraceocentrata* millions of years ago rather than hundreds of years makes the hypothesis that *P. cubensis* was brought to the Americas from its ancestral home in Africa, where it then would have to have gone extinct, less parsimonious.

The recent characterization of novel species of *Psilocybe* from Africa (Van Der Merwe et al., 2024) indicates that more diversity remains to be described there. The research landscape of *Psilocybe* has been heavily stigmatized due to the near-global government restrictions on possession of the psychoactive compounds psilocybin and psilocin. As a consequence of this long-standing impediment, research on *Psilocybe* has a rich history of cooperation between citizens and professional scientists. The study presented here was made possible by the collections and observations of numerous citizen scientists, analysis with modern phylogenetic analysis and ecological modeling, and validation against type specimens maintained in museum collections. Together, these resources provided a robust, collaborative framework that enabled the discovery of *P. ochraceocentrata* and further refined our understanding of the possible geographic origins of one of the world’s most infamous mushrooms, *P. cubensis*. More investment and less regulation on the collecting of fungi, including species and specimens predicted but not proven to contain controlled substances, is the only way to accelerate scientific discovery to gain a more complete understanding of biodiversity and its importance to human well-being before it is lost.

## ACKNOWLEDGEMENTS

The Authors would like to acknowledge Virginia Ramírez-Cruz and Laura Guzmán-Dávalos for assistance in identifying difficult-to-find specimens. Sariah VanderVeur and Toma Ipsen for assistance in sample preparation and help with technical lab duties. We would also like to thank the holding institutions who provided material, much of which is rare and irreplaceable: Instituto de Ecología (INECOL, XAL), University of Leipzig (LZ), Universidad de Guadalajara (IBUG), and Royal Botanic Gardens KEW (K). We would also like to thank Talan Moult for documenting and collecting the South African specimens of *Psilocybe ochraceocentrata* used in this study. Dr. David Minter is thanked for his thoughts and information on correct Latin designation.

## Data availability

All extracted DNA barcodes have been deposited on NCBI GenBank with corresponding accession numbers reported in Table 1. All raw genomic sequencing data has been deposited in the Short Read Archive (SRA) under Bioproject number PRJNA1159811; Biosample accession numbers are reported in Table 1. All Type specimens derived molecular data has also been provided for RefSeq designation and curation. Any code or specific script requests should be sent to the corresponding author. Supplementary data, including microscopic features, genome assembly statistics, sequence alignments, raw phylogenetic trees, and ecological and species distribution modeling outputs, can be downloaded from the DRYAD data repository at https://doi.org/10.5061/dryad.5x69p8df2.

## List of Supplementary Figures

SF1 ML tree of ITS SF2 ML tree of EF1a SF3 ML tree of RPB1 SF4 ML tree of RPB2

SF5 ENM and SDM without African occurrence data

List of Supplementary Tables:

ST1 Microscopic characteristics and measurements ST2 Genome assembly and BUSCO statistics

Supplementary data:

All occurrence data pulled from MycoPortal, including filtered sets used for ENM and SDM

## References

1. Abarenkov, K., Henrik Nilsson, R., Larsson, K., Alexander, I. J., Eberhardt, U., Erland, S., Høiland, K., Kjøller, R., Larsson, E., Pennanen, T., Sen, R., Taylor, A. F. S., Tedersoo, L., Ursing, B. M., Vrålstad, T., Liimatainen, K., Peintner, U., & Kõljalg, U. (2010). The UNITE database for molecular identification of fungi – recent updates and future perspectives. New Phytologist, 186(2), 281–285. 10.1111/j.1469-8137.2009.03160.x

2. Antón, S. C., & Swisher, Iii, C.C. (2004). Early Dispersals of *Homo* from Africa. Annual Review of Anthropology, 33(1), 271–296. 10.1146/annurev.anthro.33.070203.144024

3. Antonelli, A., Teisher, J. K., Smith, R. J., Ainsworth, A. M., Furci, G., Gaya, E., Gonçalves, S. C., Hawksworth, D. L., Larridon, I., Sessa, E. B., Simões, A. R. G., Suz, L. M., Acedo, C., Aghayeva, D. N., Agorini, A. A., Al Harthy, L. S., Bacon, K. L., Chávez Hernández, M. G., Colli Silva, M., … Williams, C. (2024). The 2030 Declaration on Scientific Plant and Fungal Collecting. *PLANTS, PEOPLE*, PLANET, ppp3.10569. 10.1002/ppp3.10569

4. Awan, A. R., Winter, J. M., Turner, D., Shaw, W. M., Suz, L. M., Bradshaw, A. J., Ellis, T., & Dentinger, B. T. M. (2018). Convergent evolution of psilocybin biosynthesis by psychedelic mushrooms. bioRxiv, 374199. 10.1101/374199

5. Belmaker, M. (2010). Early Pleistocene Faunal Connections Between Africa and Eurasia: An Ecological Perspective. In J. G. Fleagle, J. J. Shea, F. E. Grine, A. L. Baden, & R. E. Leakey (Eds.), *Out of Africa I* (pp. 183–205). Springer Netherlands. 10.1007/978-90-481-9036-2_12

6. Bengtsson Palme, J., Ryberg, M., Hartmann, M., Branco, S., Wang, Z., Godhe, A., De Wit, P., Sánchez García, M., Ebersberger, I., De Sousa, F., Amend, A., Jumpponen, A., Unterseher, M., Kristiansson, E., Abarenkov, K., Bertrand, Y. J. K., Sanli, K., Eriksson, K. M., Vik, U., … Nilsson, R. H. (2013). Improved software detection and extraction of ITS1 and ITS 2 from ribosomal ITS sequences of fungi and other eukaryotes for analysis of environmental sequencing data. Methods in Ecology and Evolution, 4(10), 914–919. 10.1111/2041-210X.12073

7. Blei, F., Dörner, S., Fricke, J., Baldeweg, F., Trottmann, F., Komor, A., Meyer, F., Hertweck, C., & Hoffmeister, D. (2020). Simultaneous Production of Psilocybin and a Cocktail of β Carboline Monoamine Oxidase Inhibitors in “Magic” Mushrooms. Chemistry – A European Journal, 26(3), 729–734. 10.1002/chem.201904363

8. Borovička, J., Noordeloos, M. E., Gryndler, M., & Oborník, M. (2011). Molecular phylogeny of Psilocybe cyanescens complex in Europe, with reference to the position of the secotioid Weraroa novae-zelandiae. Mycological Progress, 10(2), 149–155. 10.1007/s11557-010-0684-3

9. Bouckaert, R., Vaughan, T. G., Barido-Sottani, J., Duchêne, S., Fourment, M., Gavryushkina, A., Heled, J., Jones, G., Kühnert, D., De Maio, N., Matschiner, M., Mendes, F. K., Müller, N. F., Ogilvie, H. A., Du Plessis, L., Popinga, A., Rambaut, A., Rasmussen, D., Siveroni, I., … Drummond, A. J. (2019). BEAST 2.5: An advanced software platform for Bayesian evolutionary analysis. PLOS Computational Biology, 15(4), e1006650. 10.1371/journal.pcbi.1006650

10. Bradshaw, A. J., Backman, T. A., Ramírez-Cruz, V., Forrister, D. L., Winter, J. M., Guzmán- Dávalos, L., Furci, G., Stamets, P., & Dentinger, B. T. M. (2022). DNA Authentication and Chemical Analysis of *Psilocybe* Mushrooms Reveal Widespread Misdeterminations in Fungaria and Inconsistencies in Metabolites. Applied and Environmental Microbiology, 88(24), e01498–22. 10.1128/aem.01498-22

11. Bradshaw, A. J., Ramírez-Cruz, V., Awan, A. R., Furci, G., Guzmán-Dávalos, L., & Dentinger, B. T. M. (2024). Phylogenomics of the psychoactive mushroom genus *Psilocybe* and evolution of the psilocybin biosynthetic gene cluster. Proceedings of the National Academy of Sciences, 121(3), e2311245121. 10.1073/pnas.2311245121

12. Bradshaw, A., Ramírez-Cruz, V., Awan, A., Furci, G., Guzmán-Dávalos, L., & Dentinger, B. (2023). Phylogenomics of the psychoactive mushroom genus Psilocybe and evolution of the psilocybin biosynthetic gene cluster (Version 15, p. 9767543543 bytes) [Dataset]. Dryad. 10.5061/DRYAD.TMPG4F52S

13. Brown, J. L., Hill, D. J., Dolan, A. M., Carnaval, A. C., & Haywood, A. M. (2018). PaleoClim, high spatial resolution paleoclimate surfaces for global land areas. Scientific Data, 5(1), 180254. 10.1038/sdata.2018.254

14. Brownstien, M., Lazar, M., Botvinnik, A., Shevakh, C., Blakolmer, K., Lerer, L., Lifschytz, T., & Lerer, B. (2024). Striking long-term beneficial effects of single dose psilocybin and psychedelic mushroom extract in the SAPAP3 rodent model of OCD-like excessive self-grooming. Molecular Psychiatry. 10.1038/s41380-024-02786-0

15. Canan, K., Ostunii, S., Rockefeller, A., & Birkebak, J. (2024). *Psilocybe caeruleorhiza*: A new, cold weather fruiting species of psilocybin containing mushroom from the midwest in section *Aztecorum*. McIlvainea, 33. https://namyco.org/publications/mcilvainea-journal-of-american-amateur-mycology/

16. Carhart-Harris, R. L., Roseman, L., Bolstridge, M., Demetriou, L., Pannekoek, J. N., Wall, M. B., Tanner, M., Kaelen, M., McGonigle, J., Murphy, K., Leech, R., Curran, H. V., & Nutt, D. J. (2017). Psilocybin for treatment-resistant depression: fMRI-measured brain mechanisms. Scientific Reports, 7(1), 13187. 10.1038/s41598-017-13282-7

17. Castro Jauregui, O. S., Ramírez-Cruz, V., Bradshaw, A. J., Cortés-Pérez, A., & Guzmán-Dávalos, L. (2022). Los hongos sagrados del género Psilocybe en Jalisco. Nubes y Ciencia, 12, 14–21.

18. Cerling, T. E. (1992). Development of grasslands and savannas in East Africa during the Neogene. *Palaeogeography, Palaeoclimatology*, Palaeoecology, 97(3), 241–247. 10.1016/0031-0182(92)90211-M

19. Cerling, T. E., Bowman, J. R., & O’Neil, J. R. (1988). An isotopic study of a fluvial-lacustrine sequence: The Plio-Pleistocene koobi fora sequence, East Africa. Palaeogeography, Palaeoclimatology, Palaeoecology, 63(4), 335–356. 10.1016/0031-0182(88)90104-6

20. Chen, S., Zhou, Y., Chen, Y., & Gu, J. (2018). fastp: An ultra-fast all-in-one FASTQ preprocessor. Bioinformatics, 34(17), i884–i890. 10.1093/bioinformatics/bty560

21. Crous, P. W., Rong, I. H., Wood, A., Lee, S., Glen, H., Botha, W., Slippers, B., De Beer, W. Z., Wingfield, M. J., & Hawksworth, D. L. (2006). How many species of fungi are there at the tip of Africa? Studies in Mycology, 55, 13–33. 10.3114/sim.55.1.13

22. Daniel, J., & Haberman, M. (2017). Clinical potential of psilocybin as a treatment for mental health conditions. Mental Health Clinician, 7(1), 24–28. 10.9740/mhc.2017.01.024

23. Dantas, M. A. T., Campbell, S. C., & McDonald, H. G. (2023). Paleoecological inferences about the Late Quaternary giant sloths. Journal of Mammalian Evolution, 30(4), 891–905. 10.1007/s10914-023-09681-5

24. Dennell, R., & Roebroeks, W. (2005). An Asian perspective on early human dispersal from Africa. Nature, 438(7071), 1099–1104. 10.1038/nature04259

25. Dentinger, B. T. M., Margaritescu, S., & Moncalvo, J. (2010). Rapid and reliable high throughput methods of DNA extraction for use in barcoding and molecular systematics of mushrooms. Molecular Ecology Resources, 10(4), 628–633. 10.1111/j.1755-0998.2009.02825.x

26. Dolan, A. M., Haywood, A. M., Hunter, S. J., Tindall, J. C., Dowsett, H. J., Hill, D. J., & Pickering, S. J. (2015). Modelling the enigmatic Late Pliocene Glacial Event—Marine Isotope Stage M2. Global and Planetary Change, 128, 47–60. 10.1016/j.gloplacha.2015.02.001

27. Drummond, A. J., & Rambaut, A. (2007). BEAST: Bayesian evolutionary analysis by sampling trees. BMC Evolutionary Biology, 7(1), 214. 10.1186/1471-2148-7-214

28. Edwards, E. J., Osborne, C. P., Strömberg, C. A. E., Smith, S. A., C Grasses Consortium, Bond, W. J., Christin, P.-A., Cousins, A. B., Duvall, M. R., Fox, D. L., Freckleton, R. P., Ghannoum, O., Hartwell, J., Huang, Y., Janis, C. M., Keeley, J. E., Kellogg, E. A., Knapp, A. K., Leakey, A. D. B., … Tipple, B. (2010). The Origins of C _4_ Grasslands: Integrating Evolutionary and Ecosystem Science. Science, 328(5978), 587–591. 10.1126/science.1177216

29. Fick, S. E., & Hijmans, R. J. (2017). WorldClim 2: New 1 km spatial resolution climate surfaces for global land areas. International Journal of Climatology, 37(12), 4302–4315. 10.1002/joc.5086

30. Fricke, J., Blei, F., & Hoffmeister, D. (2017). Enzymatic Synthesis of Psilocybin. Angewandte Chemie International Edition, 56(40), 12352–12355. 10.1002/anie.201705489

31. Froese, D., Stiller, M., Heintzman, P. D., Reyes, A. V., Zazula, G. D., Soares, A. E. R., Meyer, M., Hall, E., Jensen, B. J. L., Arnold, L. J., MacPhee, R. D. E., & Shapiro, B. (2017). Fossil and genomic evidence constrains the timing of bison arrival in North America. Proceedings of the National Academy of Sciences, 114(13), 3457–3462. 10.1073/pnas.1620754114

32. Froese, T., Guzmán, G., & Guzmán-Dávalos, L. (2016). On the Origin of the Genus Psilocybe and Its Potential Ritual Use in Ancient Africa and Europe1. Economic Botany, 70(2), 103–114. 10.1007/s12231-016-9342-2

33. Gandy, S., Forstmann, M., Carhart-Harris, R. L., Timmermann, C., Luke, D., & Watts, R. (2020). The potential synergistic effects between psychedelic administration and nature contact for the improvement of mental health. Health Psychology Open, 7(2), 205510292097812. 10.1177/2055102920978123

34. Gates, C. C., Freese, C. H., Gogan, P. J., & Kotzman, M. (2010). American bison: Status survey and conservation guidelines 2010. IUCN. https://books.google.com/books?hl=en&lr=&id=koUrGx-i2ucC&oi=fnd&pg=PR11&dq=C.+C.+Gates,+C.+H.+Freese,+P.+J.+Gogan,+M.+Kotzman,+American+Bison:+Status+Survey+and+Conservation+Guidelines+2010+(IUCN,+Gland,+Switzerland,+2010).&ots=VYEfbm1jA_&sig=FAszmMQVic5vjaMvgjnjC7MK2EE

35. Guzmán, G. (1983). The genus Psilocybe: A systematic revision of the known species including the history, distribution, and chemistry of the hallucinogenic species. J. Cramer.

36. Guzmán, G. (1995). Supplement to the monograph of the genus Psilocybe.

37. Guzman, G. (2005). Species Diversity of the Genus Psilocybe (Basidiomycotina, Agaricales, Strophariaceae) in the World Mycobiota, with Special Attention to Hallucinogenic Properties. International Journal of Medicinal Mushrooms, 7(1–2), 305–332. 10.1615/IntJMedMushr.v7.i12.280

38. Guzmán, G. (2008). Hallucinogenic Mushrooms in Mexico: An Overview. Economic Botany, 62(3), 404–412. 10.1007/s12231-008-9033-8

39. Guzmán, G. (2014). Psilocybe s. Str. (Agaricales, Strophariaceae) in Africa with description of a new species from the Congo. Sydowia An International Journal of Mycology, 66, 43–53. 10.12905/0380.sydowia66(1)2014-0043

40. Guzman, G., Cortes-Perez, A., & Ramirez-Guillen, F. (2013). The Japanese Hallucinogenic Mushrooms Psilocybe and a New Synonym of P. subcaerulipes with Three Asiatic Species Belong to Section Zapotecorum (Higher Basidiomycetes). International Journal of Medicinal Mushrooms, 15(6), 607–615. 10.1615/IntJMedMushr.v15.i6.90

42. Handika, H., & Esselstyn, J. (2022). *SEGUL: An ultrafast, memory-efficient alignment manipulation and summary tool for phylogenomics* [Preprint]. Preprints. 10.22541/au.165167823.30911834/v1

43. Handika, H., & Esselstyn, J. A. (2024). SEGUL: Ultrafast, memory efficient and mobile friendly software for manipulating and summarizing phylogenomic datasets. *Molecular Ecology Resources*, e13964. 10.1111/1755-0998.13964

44. Hanotte, O., Bradley, D. G., Ochieng, J. W., Verjee, Y., Hill, E. W., & Rege, J. E. O. (2002). African Pastoralism: Genetic Imprints of Origins and Migrations. Science, 296(5566), 336–339. 10.1126/science.1069878

45. Hill, D. J. (2015). The non-analogue nature of Pliocene temperature gradients. Earth and Planetary Science Letters, 425, 232–241. 10.1016/j.epsl.2015.05.044

46. Johnson, M. W., & Griffiths, R. R. (2017). Potential Therapeutic Effects of Psilocybin. Neurotherapeutics, 14(3), 734–740. 10.1007/s13311-017-0542-y

47. Johnston, P. }, & Buchanan, P. K. (1995). The genus *Psilocybe* (Agaricales) in New Zealand. New Zealand Journal of Botany, 33(3), 379–388. 10.1080/0028825X.1995.10412964

48. Kahle, D., & Wickham, H. (2013). ggmap: Spatial Visualization with ggplot2. The R Journal, 5(1), 144. 10.32614/RJ-2013-014

49. Kalyaanamoorthy, S., Minh, B. Q., Wong, T. K. F., von Haeseler, A., & Jermiin, L. S. (2017). ModelFinder: Fast model selection for accurate phylogenetic estimates. Nature Methods, 14(6), 587–589. 10.1038/nmeth.4285

50. Karger, D. N., Conrad, O., Böhner, J., Kawohl, T., Kreft, H., Soria-Auza, R. W., Zimmermann, N. E., Linder, H. P., & Kessler, M. (2017). Climatologies at high resolution for the earth’s land surface areas. Scientific Data, 4(1), 170122. 10.1038/sdata.2017.122

51. Katoh, K. (2002). MAFFT: A novel method for rapid multiple sequence alignment based on fast

52. Fourier transform. Nucleic Acids Research, 30(14), 3059–3066. 10.1093/nar/gkf436

53. Lerer, E., Botvinnik, A., Shahar, O., Grad, M., Blakolmer, K., Shomron, N., Lotan, A., Lerer, B., & Lifschytz, T. (2024). Effects of psilocybin, psychedelic mushroom extract and 5- hydroxytryptophan on brain immediate early gene expression: Interaction with serotonergic receptor modulators. Frontiers in Pharmacology, 15, 1391412. 10.3389/fphar.2024.1391412

54. Ma, T., Feng, Y., Lin, X.-F., Karunarathna, S. C., Ding, W.-F., & Hyde, K. D. (2014). Psilocybe chuxiongensis, a new bluing species from subtropical China. Phytotaxa, 156(4), 211. 10.11646/phytotaxa.156.4.3

55. Matsushima, Y., Shirota, O., Kikura-Hanajiri, R., Goda, Y., & Eguchi, F. (2009). Effects of *Psilocybe argentipes* on Marble-Burying Behavior in Mice. Bioscience, Biotechnology, and Biochemistry, 73(8), 1866–1868. 10.1271/bbb.90095

56. McTaggart, A. R., McLaughlin, S., Slot, J. C., McKernan, K., Appleyard, C., Bartlett, T. L., Weinert, M., Barlow, C., Warne, L. N., Shuey, L. S., Drenth, A., & James, T. Y. (2023). Domestication through clandestine cultivation constrained genetic diversity in magic mushrooms relative to naturalized populations. Current Biology, 33(23), 5147–5159.e7. 10.1016/j.cub.2023.10.059

57. Miller, A. N., & Bates, S. T. (2017). The Mycology Collections Portal (MyCoPortal). IMA Fungus, 8(2), A65–A66. 10.1007/BF03449464

58. Minh, B. Q., Nguyen, M. A. T., & von Haeseler, A. (2013). Ultrafast Approximation for Phylogenetic Bootstrap. Molecular Biology and Evolution, 30(5), 1188–1195. 10.1093/molbev/mst024

59. Musshoff, F., Madea, B., & Beike, J. (2000). Hallucinogenic mushrooms on the German market—Simple instructions for examination and identification. Forensic Science International, 113(1–3), 389–395. 10.1016/S0379-0738(00)00211-5

60. Nguyen, L.-T., Schmidt, H. A., von Haeseler, A., & Minh, B. Q. (2015). IQ-TREE: A Fast and Effective Stochastic Algorithm for Estimating Maximum-Likelihood Phylogenies.

61. Molecular Biology and Evolution, 32(1), 268–274. 10.1093/molbev/msu300

62. Nurk, S., Meleshko, D., Korobeynikov, A., & Pevzner, P. A. (2017). metaSPAdes: A new versatile metagenomic assembler. Genome Research, 27(5), 824–834. 10.1101/gr.213959.116

63. Ostunii, S., Rockefeller, A., Jacobs, J., & Birkebak, J. (2024). *Psilocybe niveotropicalis*: A new species of psilocybin containing mushroom from South Florida. McIlvainea, 33. https://namyco.org/publications/mcilvainea-journal-of-american-amateur-mycology/

64. Pebesma, E. (2018). Simple Features for R: Standardized Support for Spatial Vector Data. The R Journal, 10(1), 439. 10.32614/RJ-2018-009

65. Picker, J., & Rickards, R. (1970). The occurrence of the psychotomimetic agent psilocybin in an Australian agaric, Psilocybe subaeruginosa. Australian Journal of Chemistry, 23(4), 853. 10.1071/CH9700853

66. Piepenbring, M., Maciá-Vicente, J. G., Codjia, J. E. I., Glatthorn, C., Kirk, P., Meswaet, Y., Minter, D., Olou, B. A., Reschke, K., Schmidt, M., & Yorou, N. S. (2020). Mapping mycological ignorance – checklists and diversity patterns of fungi known for West Africa. IMA Fungus, 11(1), 13. 10.1186/s43008-020-00034-y

67. Pitt, D., Sevane, N., Nicolazzi, E. L., MacHugh, D. E., Park, S. D. E., Colli, L., Martinez, R., Bruford, M. W., & Orozco terWengel, P. (2019). Domestication of cattle: Two or three events? Evolutionary Applications, 12(1), 123–136. 10.1111/eva.12674

68. Prjibelski, A., Antipov, D., Meleshko, D., Lapidus, A., & Korobeynikov, A. (2020). Using SPAdes De Novo Assembler. Current Protocols in Bioinformatics, 70(1), e102. 10.1002/cpbi.102

69. Rambaut, A., Drummond, A. J., Xie, D., Baele, G., & Suchard, M. A. (2018). Posterior Summarization in Bayesian Phylogenetics Using Tracer 1.7. Systematic Biology, 67(5), 901–904. 10.1093/sysbio/syy032

70. Ramírez-Cruz, V., Guzmán, G., Villalobos-Arámbula, A. R., Rodríguez, A., Matheny, P. B., Sánchez-García, M., & Guzmán-Dávalos, L. (2013). Phylogenetic inference and trait evolution of the psychedelic mushroom genus *Psilocybe* sensu lato (Agaricales). Botany, 91(9), 573–591. 10.1139/cjb-2013-0070

71. Rayner, R. W. (1970). A mycological colour chart. Commonwealth Mycological Institute ; British Mycological Society.

72. Ripple, W. J., Newsome, T. M., Wolf, C., Dirzo, R., Everatt, K. T., Galetti, M., Hayward, M. W., Kerley, G. I. H., Levi, T., Lindsey, P. A., Macdonald, D. W., Malhi, Y., Painter, L. E., Sandom, C. J., Terborgh, J., & Van Valkenburgh, B. (2015). Collapse of the world’s largest herbivores. Science Advances, 1(4), e1400103. 10.1126/sciadv.1400103

73. Sayers, E. W., Cavanaugh, M., Clark, K., Ostell, J., Pruitt, K. D., & Karsch-Mizrachi, I. (2019).

74. GenBank. Nucleic Acids Research,gkz956. 10.1093/nar/gkz956

75. Schoch, C. L., Seifert, K. A., Huhndorf, S., Robert, V., Spouge, J. L., Levesque, C. A., Chen, W., Fungal Barcoding Consortium, Fungal Barcoding Consortium Author List, Bolchacova, E., Voigt, K., Crous, P. W., Miller, A. N., Wingfield, M. J., Aime, M. C., An, K.-D., Bai, F.- Y., Barreto, R. W., Begerow, D., … Schindel, D. (2012). Nuclear ribosomal internal transcribed spacer (ITS) region as a universal DNA barcode marker for Fungi.Proceedings of the National Academy of Sciences, 109(16), 6241–6246. 10.1073/pnas.1117018109

76. Shahar, O., Botvinnik, A., Shwartz, A., Lerer, E., Golding, P., Buko, A., Hamid, E., Kahn, D., Guralnick, M., Blakolmer, K., Wolf, G., Lotan, A., Lerer, L., Lerer, B., & Lifschytz, T. (2024). Effect of chemically synthesized psilocybin and psychedelic mushroom extract on molecular and metabolic profiles in mouse brain. Molecular Psychiatry, 29(7), 2059–2073. 10.1038/s41380-024-02477-w

77. Shlemov, A., & Korobeynikov, A. (2019). PathRacer: Racing profile HMM paths on assembly graph. 10.1101/562579

78. Simão, F. A., Waterhouse, R. M., Ioannidis, P., Kriventseva, E. V., & Zdobnov, E. M. (2015). BUSCO: Assessing genome assembly and annotation completeness with single-copy orthologs. Bioinformatics, 31(19), 3210–3212. 10.1093/bioinformatics/btv351

79. Sluyter, A. (2023). Cattle in Latin American History. In A. Sluyter, Oxford Research Encyclopedia of Latin American History. Oxford University Press. 10.1093/acrefore/9780199366439.013.1153

80. Thomas, K., & Manimohan, P. (2003). The genus Agrocybe in Kerala State, India. Mycotaxon. https://www.semanticscholar.org/paper/The-genus-Agrocybe-in-Kerala-State%2C-India.-Thomas-Manimohan/919ec55a40a6313691253205ee5104b88a3dc96e

81. Thomas, K., Manimohan, P., Guzmán, G., Tapia, F., & Ramirez-Guillen, F. (2002). The genus Psilocybe in Kerala State, India. Mycotaxon -Ithaca Ny-, 83, 195–207.

82. Tsakem, B., Tchamgoue, J., Kinge, R. T., Tiani, G. L. M., Teponno, R. B., & Kouam, S. F. (2024). Diversity of African fungi, chemical constituents and biological activities. Fitoterapia, 178, 106154. 10.1016/j.fitote.2024.106154

83. Van Court, R. C., Wiseman, M. S., Meyer, K. W., Ballhorn, D. J., Amses, K. R., Slot, J. C., Dentinger, B. T. M., Garibay-Orijel, R., & Uehling, J. K. (2022). Diversity, biology, and history of psilocybin-containing fungi: Suggestions for research and technological development. Fungal Biology, 126(4), 308–319. 10.1016/j.funbio.2022.01.003

84. Van Der Merwe, B., Rockefeller, A., Kilian, A., Clark, C., Sethathi, M., Moult, T., & Jacobs, K. (2024). A description of two novel *Psilocybe* species from southern Africa and some notes on African traditional hallucinogenic mushroom use. Mycologia, 1–14. 10.1080/00275514.2024.2363137

85. White, T., Bruns, T., Lee, S., Taylor, J., Innis, M., Gelfand, D., & Sninsky, J. (1990). Amplification and Direct Sequencing of Fungal Ribosomal RNA Genes for Phylogenetics. In Pcr Protocols: A Guide to Methods and Applications, (Vol. 31, pp. 315–322).

86. Wickham, H. (2016). ggplot2: Elegant Graphics for Data Analysis (2nd ed. 2016). Springer International Publishing : Imprint: Springer. 10.1007/978-3-319-24277-4

87. Zeuner, F. E. (1963). A History of Domesticated Animals. Hutchinson. Zhu, Z., Dennell, R., Huang, W., Wu, Y., Qiu, S., Yang, S., Rao, Z., Hou, Y., Xie, J., Han, J., & Ouyang, T. (2018). Hominin occupation of the Chinese Loess Plateau since about 2.1 million years ago. Nature, 559(7715), 608–612. 10.1038/s41586-018-0299-4

88. Zhuk, O., Jasicka-Misiak, I., Poliwoda, A., Kazakova, A., Godovan, V., Halama, M., & Wieczorek, P. (2015). Research on Acute Toxicity and the Behavioral Effects of Methanolic Extract from Psilocybin Mushrooms and Psilocin in Mice. Toxins, 7(4), 1018– 1029. 10.3390/toxins7041018

