## Supplementary material for "Discovery of the closest free-living relative of the domesticated “magic mushroom” *Psilocybe cubensis* in Africa": Table 1: Psilocybe_ochraceocentrat_voucher_table.html

Specimen Voucher Table

| Sample name | Voucher designation | Determination | Locality | Type status | Collection year | Type authorship | Holding institution | Herbarium Code | Catalog number | Accession\|ITS | Accession\|EF1a | Accession\|RPB1 | Accession\|RPB2 | SRA | BioSample |
| --- | --- | --- | --- | --- | --- | --- | --- | --- | --- | --- | --- | --- | --- | --- | --- |
| Psilocybe\_sp\_Africa\_Zimbabwe\_CS-3006 | Psilocybe sp. | Psilocybe ochraceocentrata sp. nov. | Zimbabwe (Matabeleland South Province) | Holotype | 2013 | This publication | Natural History Museum of Zimbabwe | BUL | BUL8013 | PQ315824 | PQ317397 | PQ317404 | PQ317412 | SRR30663520 | SAMN43585389 |
| Psilocybe\_Africa\_Zimbabwe\_CS-5783 | Psilocybe sp. | Psilocybe ochraceocentrata sp. nov. | Zimbabwe (Matabeleland South Province) | N/A | 2018 | This publication | Private Fungarium of C. Sharp | N/A | CS-5783 | PQ315825 | PQ317398 | PQ317405 | PQ317413 | SRR30663519 | SAMN43585390 |
| Psilocybe\_sp\_Africa\_Zimbabwe\_CS-3309 | Psilocybe sp. | Psilocybe ochraceocentrata sp. nov. | Zimbabwe (Mashonaland Central Province) | N/A | 2013 | This publication | Private Fungarium of C. Sharp | N/A | CS-3309 | PQ315826 | PQ317399 | PQ317406 | PQ317414 | SRR30663518 | SAMN43585390 |
| T3 | Psilocybe ochraceocentrata | Psilocybe ochraceocentrata sp. nov. | South Africa (Kwa-Zulu Natal province) | N/A | 2022 | This publication | Stellenbosch University | N/A | Psilocybe ochraceocentrata -T1 | PP959211 | N/A | N/A | N/A | N/A | N/A |
| T1 | Psilocybe ochraceocentrata | Psilocybe ochraceocentrata sp. nov. | South Africa (Kwa-Zulu Natal province) | N/A | 2022 | This publication | Stellenbosch University | N/A | Psilocybe ochraceocentrata -T3 | PP959212 | N/A | N/A | N/A | N/A | N/A |
| Harding | Psilocybe ochraceocentrata | Psilocybe ochraceocentrata sp. nov. | South Africa (Kwa-Zulu Natal province) | N/A | 2022 | This publication | Stellenbosch University | N/A | Psilocybe ochraceocentrata- Harding | PP959213 | N/A | N/A | N/A | N/A | N/A |
| Stropharia\_cubenesis\_NY-761563\_Holotype | Stropharia cubensis | Psilocybe cubensis (Earle) Singer | Cuba | Holotype | 1906 | Sydowia 2(1-6): 37 (1948) | New York Botanical Garden | NY | NY-761563 | PQ315827 | Not recovered | PQ317407 | PQ317415 | SRR30663517 | SAMN43585392 |
| Psilocybe\_jaliscana\_IBUG-13710\_Isotype | Psilocybe jaliscana | Psilocybe cubensis (Earle) Singer | Meico, Jalisco | Isotype | 2000 | Docums Mycol. 29(no. 116): 46 (2000) | Universidad de Guadalajara | IBUG | IBUG-13710 | PQ315828 | PQ317400 | PQ317408 | PQ317416 | SRR30663516 | SAMN43585393 |
| Psilocybe\_natalensis\_Lz\_Psi\_nat\_reseq\_HOLOTYPE | Psilocybe natalensis | Psilocybe natalensis Gartz, D.A. Reid, M.T. Sm. & Eicker | KwaZulu-Natal | Holotype | 1994 | Integration 6: 29 (1995) | University of Leipzig | LZ | 6266 | PQ315829 | PQ317401 | PQ317409 | PQ317417 | SRR30663515 | SAMN43585394 |
| Psilocybe\_wayanadensis\_XAL-Thomas-T-320a\_Holotype | Psilocybe wayanadensis | Psilocybe wayanadensis K.A. Thomas, Manim. & Guzmán, | India, Kerala | Holotype | 2002 | Mycotaxon 83: 198 (2002) | Instituto de Ecología, A.C. | XAL | Not Recorded | PQ315830 | PQ317402 | PQ317410 | PQ317418 | SRR30663514 | SAMN43585395 |
| Psilocybe\_keralensis\_XAL-T319a\_Isotype | Psilocybe keralensis | Psilocybe keralensis K.A. Thomas, Manim. & Guzmán | India, Kerala | Isotype | 2002 | Mycotaxon 83: 196 (2002) | Instituto de Ecología, A.C. | XAL | Not Recorded | PQ315831 | PQ317403 | PQ317411 | PQ317419 | SRR30663513 | SAMN43585396 |
| Stropharia\_aquamarina\_Holotype\_KM-160812 | Stropharia aquamarina | Psilocybe aquamarina (Pegler) Guzmán | Kenya | Holotype | 1977 | Biblthca Mycol. 159: 109 (1995) | Royal Botanic Gardens, KEW | K(M) | KM-160812 | PQ330245 | PQ435231 | PQ435232 | PQ435233 | SRR30690310 | SAMN43784757 |
