## Supplemental Figures1-5 for "Discovery of the closest free-living relative of the domesticated “magic mushroom” *Psilocybe cubensis* in Africa"

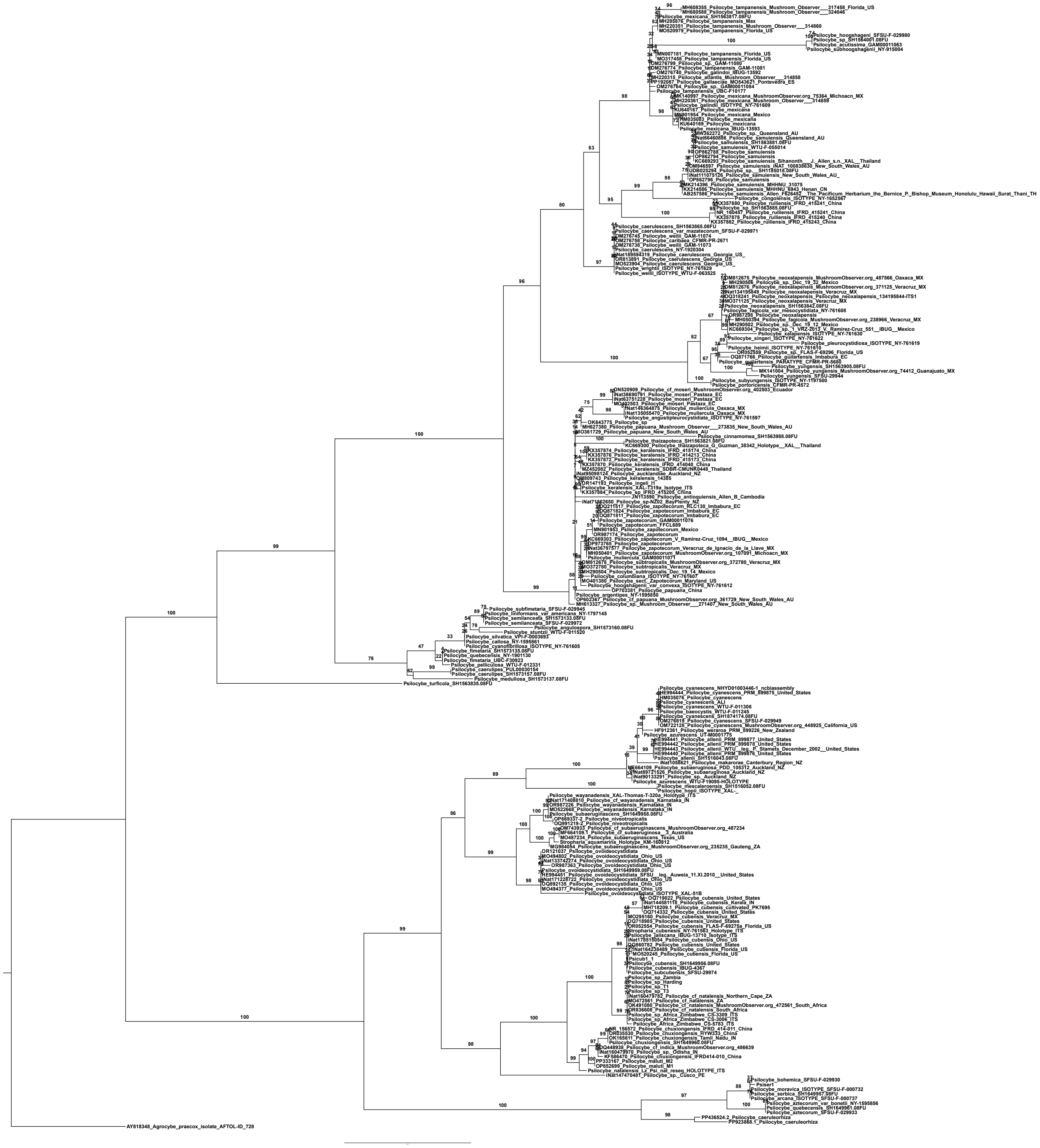

Supplementary Figure 1.
Maximum likelihood phylogram generated with currated ITS sequences from NCBI GenBank, UNITE, and previously published studies.Branch labels percent ultrafast bootstraps (BS) (n=1000). Tree is rooted *Agrocybe praexcox* AFTOL-ID 728.

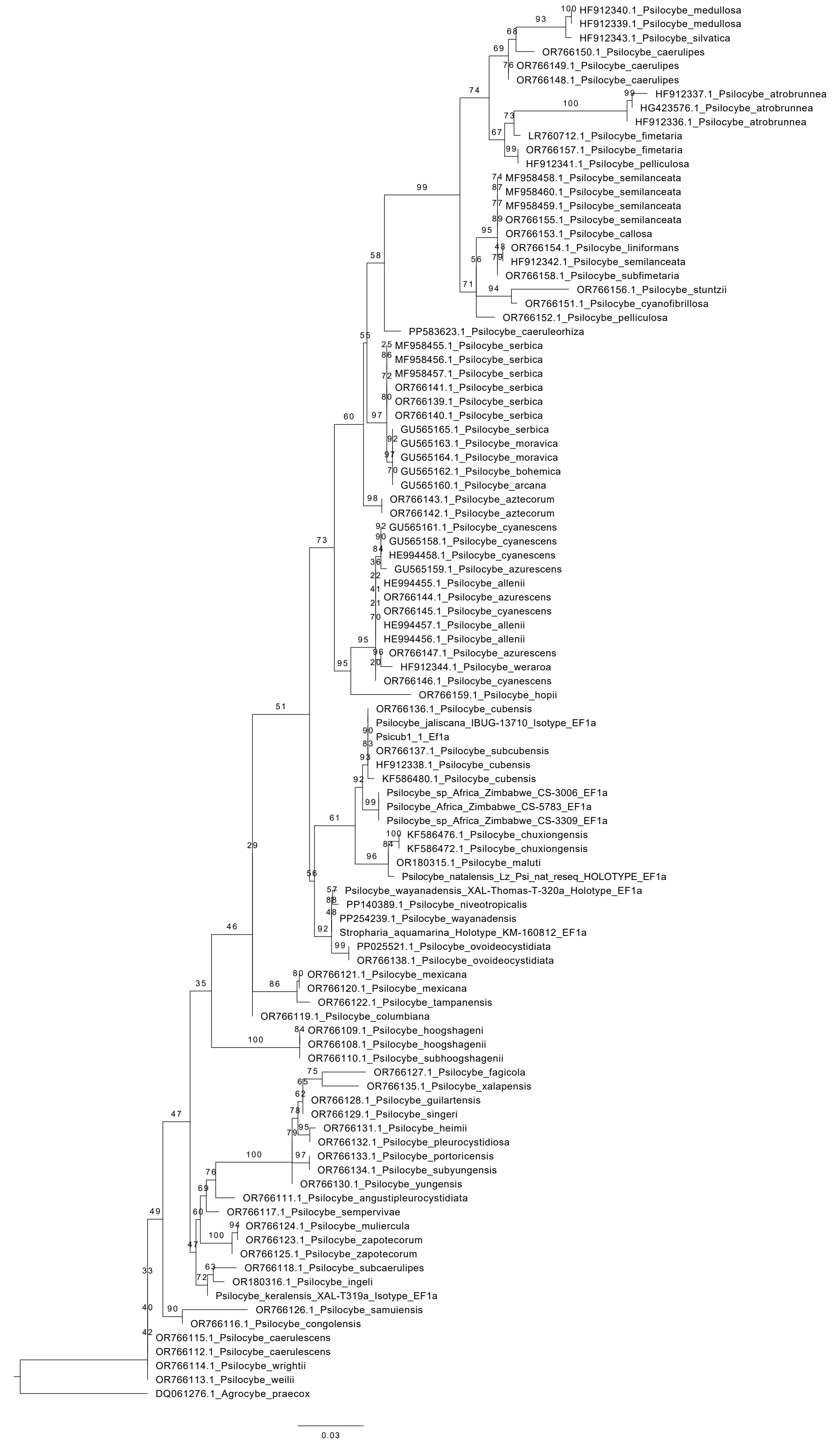


Supplementary Figure 2.
Maximum likelihood phylogram generated with publically available EF1a sequences from NCBI GenBank. Branch labels percent ultrafast bootstraps (BS) (n=1000).
Tree is rooted *Agrocybe praexcox* DQ061276.1.

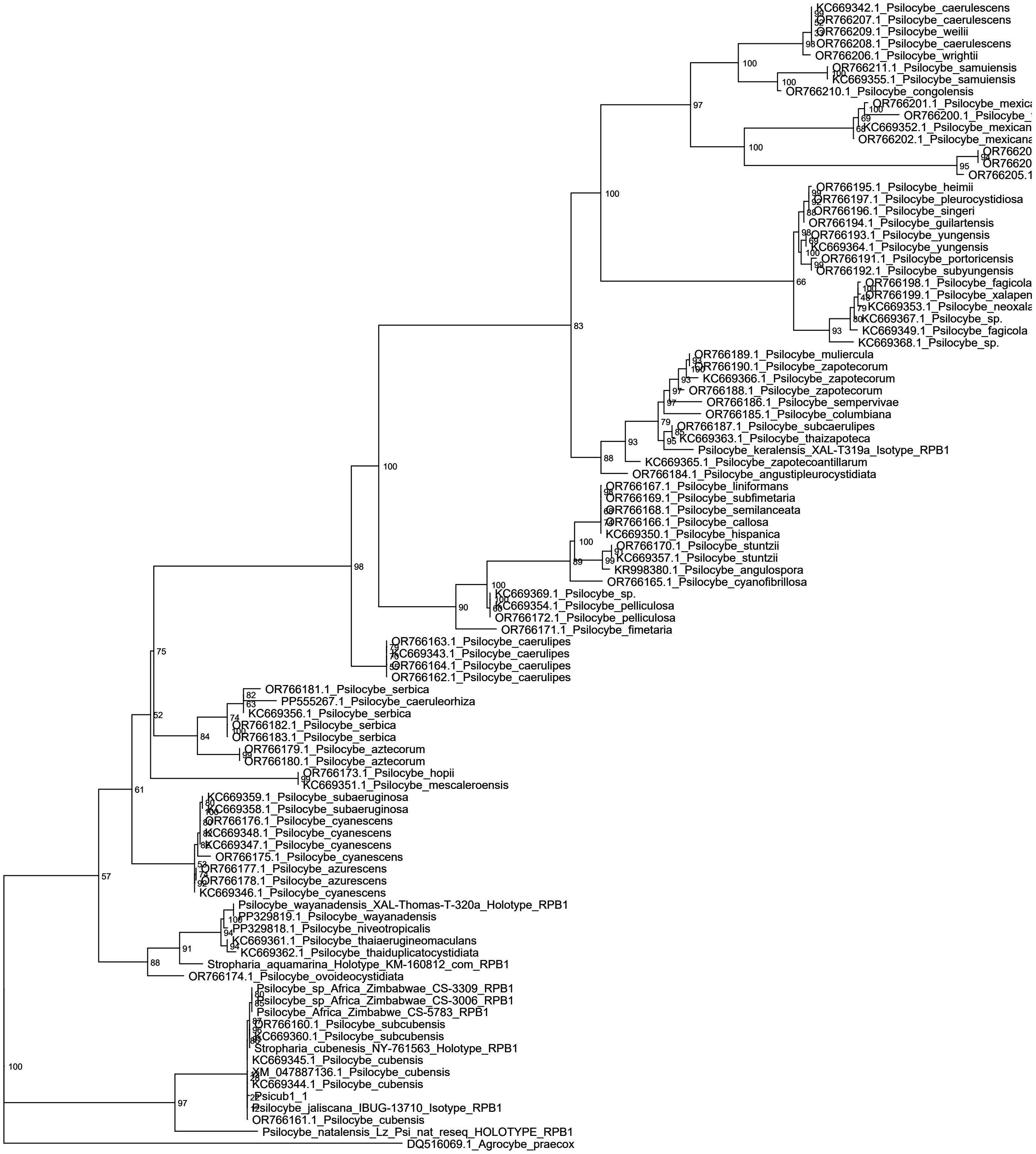


Supplementary Figure 3.
Maximum likelihood phylogram generated with publically available RPB1 sequences from NCBI GenBank. Branch labels percent ultrafast bootstraps (BS) (n=1000).
Tree is rooted *Agrocybe praexcox* DQ516069.1.

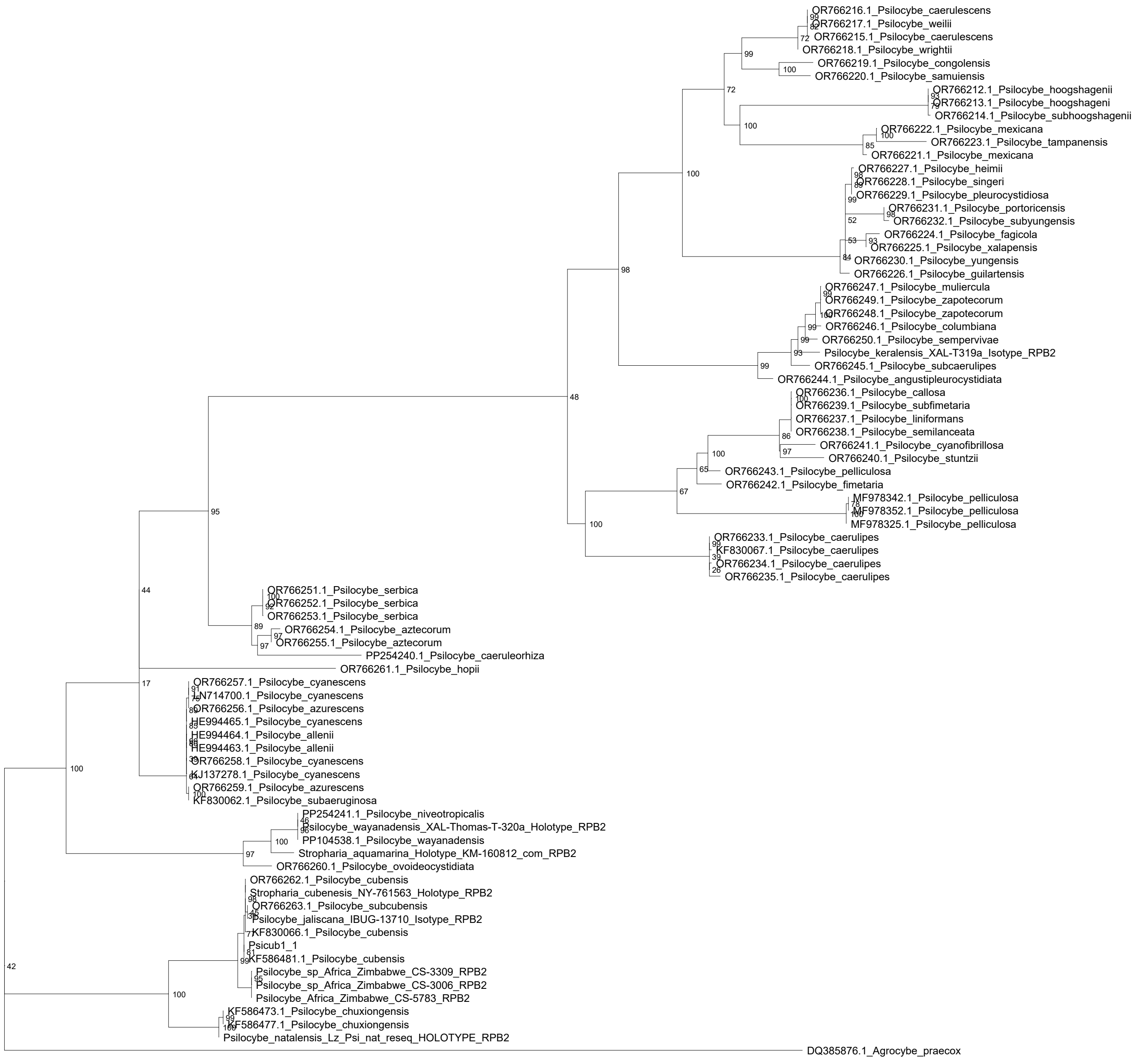


Supplementary Figure 4.
Maximum likelihood phylogram generated with publically available RPB2 sequences from NCBI GenBank. Branch labels percent ultrafast bootstraps (BS) (n=1000).
Tree is rooted *Agrocybe praexcox* DQ385876.1.

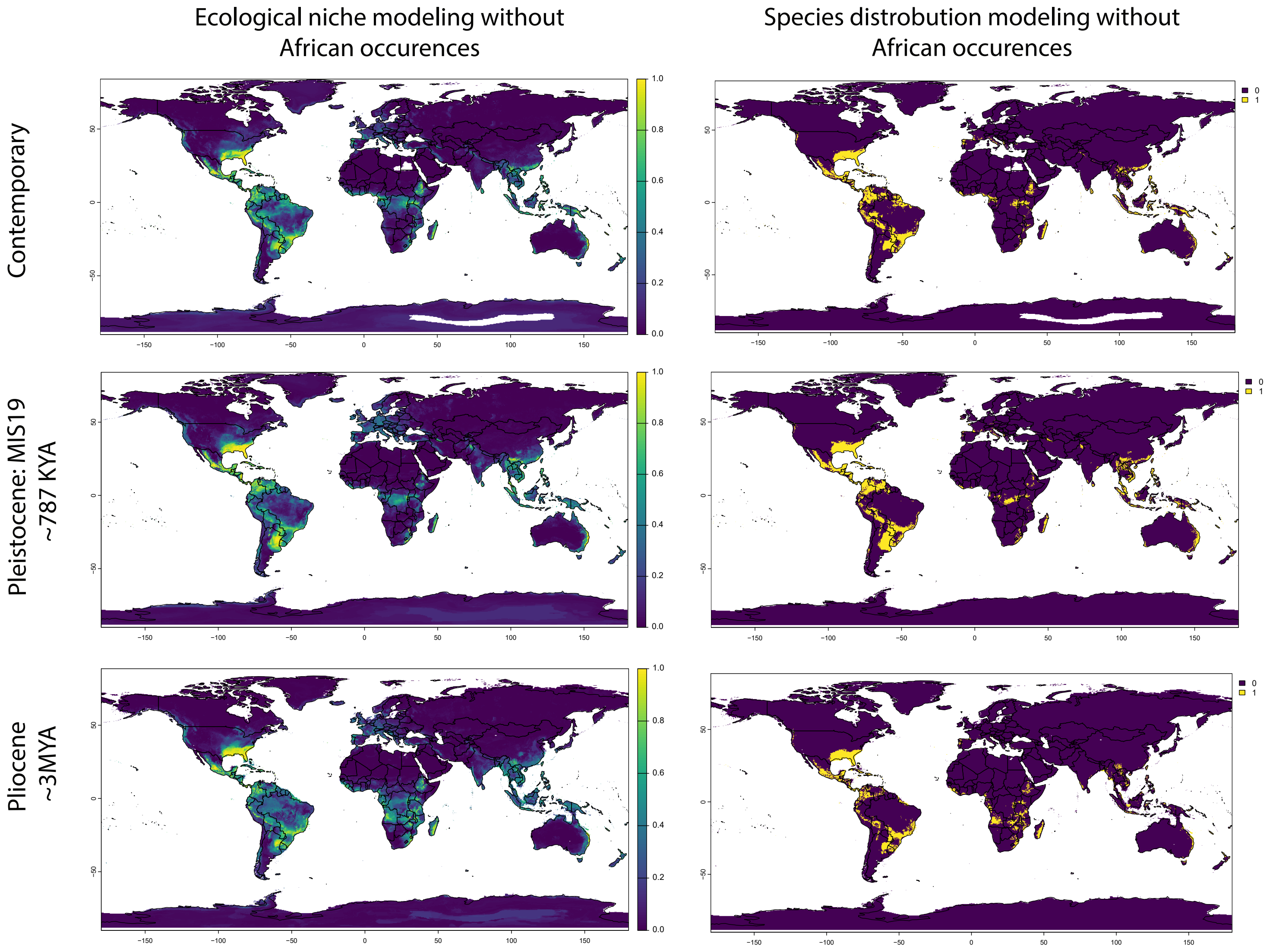


Supplementary Figure 5.
Ecological niche and Species distribution modeling of *Psilocybe Cubensis* throughout time.
ENM and SDM of *P. cubensis*, excluding African occurrence data from public sources across three major time scales: Top; Contemporary (Modern day), Middle; Pleistocene MS19 (~787KYA). Bottom; Pliocene (~3MYA). Left: ENM with distribution likelihood indicated as a heat gradient from purple (0%) to yellow (100%). Right: SDM with predicted species presence as present (1, yellow)  or absent (0, purple).
